# Cross-species epigenetic regulation of nucleus accumbens *KCNN3* transcript variants by excessive ethanol drinking and dependence

**DOI:** 10.1101/713826

**Authors:** Rita Cervera-Juanes, Audrey E. Padula, Larry J. Wilhem, Byung Park, Kathleen A. Grant, Betsy Ferguson, Patrick J. Mulholland

## Abstract

The underlying genetic and epigenetic mechanisms driving functional adaptations in neuronal excitability and excessive alcohol intake are poorly understood. Given that small-conductance Ca^2+^-activated K^+^ (K_Ca_2 or SK) channels encoded by the *KCNN* family of genes have emerged from preclinical studies as a crucial target that contributes to heavy drinking and alcohol-induced functional neuroadaptations, we performed a cross-species analysis of *KCNN3* methylation, gene expression, and polymorphisms of alcohol-drinking monkeys and alcohol dependent mice. Because of the alternative promoters in *KCNN3*, we analyzed expression of the different transcript variants that when translated influence surface trafficking and function of K_Ca_2 channels. In heavy drinking rhesus macaques and alcohol dependent C57BL/6J mice, bisulfite sequencing analysis of the nucleus accumbens revealed a differentially methylated region in exon 1A of *KCNN3* that overlaps with a predicted promoter sequence. The hypermethylation of *KCNN3* in monkey and mouse accumbens paralleled an increase in expression of alternative transcript variants that encode apamin-insensitive and dominant-negative K_Ca_2 channel isoforms. A polymorphic repeat in macaque *KCNN3* encoded by exon 1 did not correlate with alcohol drinking. At the protein level, K_Ca_2.3 channel expression in the accumbens was significantly reduced in very heavy drinking monkeys. Together, our cross-species findings on epigenetic dysregulation of *KCNN3* by heavy alcohol drinking and dependence represent a complex mechanism that utilizes alternative promoters to impact firing of accumbens neurons. Thus, these results provide support for hypermethylation of *KCNN3* by excessive alcohol drinking as a possible key molecular mechanism underlying harmful alcohol intake and alcohol use disorder.

## INTRODUCTION

Alcohol (ethanol) use disorder (AUD) is a devastating brain disease driven by complex interactions between genetic, epigenetic, and environmental factors. Excessive ethanol drinking is a leading cause of preventable death and was estimated to cost the U.S. $249 billion in 2010, $100 billion of which was paid for by the government (1, 2). There are genetic factors that increase the propensity for risky drinking (3, 4), and genetic background influences the efficacy of treatment options for individuals with AUD (5). In addition, prolonged excessive ethanol intake produces neuroadaptations in projection neurons and neural circuits that are proposed to sustain heavy drinking (6–8). Although there are three FDA-approved pharmacotherapies for treating AUD, only one (i.e., naltrexone) targets genetic variation in individuals with AUD and the others do not address adaptations that facilitate altered firing properties in neurons. While pharmacologically targeting known polymorphisms can reduce relapse rates in a subpopulation of individuals with AUD (5, 9), there have been mixed results that suggest a critical need for further investigation of the genetic factors and neuroadaptations that contribute to excessive ethanol intake.

Small-conductance Ca^2+^-activated K^+^ (K_Ca_2 or SK) channels have emerged from preclinical pharmacogenetic studies as a target for treating AUD (6, 10–14). K_Ca_2.3 channels are encoded by the *KCNN3* gene and are enriched in nucleus accumbens core (NAcC) medium spiny neurons (MSNs) and substantia nigra dopamine neurons where they control neuronal firing patterns (15). In previous integrative functional genomics studies, we reported that cross-species ethanol-related QTLs contain *KCNN3*, and NAcC *Kcnn3* transcript levels negatively correlated with voluntary drinking in genetically diverse BXD strains of mice (13, 14). Moreover, excessive differentiation in the promoter region of *Kcnn3* associated with ethanol preference in selectively bred rat lines (16). Induction of ethanol dependence and excessive drinking increased intrinsic excitability and reduced K_Ca_2 channel function and K_Ca_2.3 channel protein expression in the NAcC of rodents(11, 13, 17). Blocking K_Ca_2 channels in the NAcC with apamin increased voluntary ethanol drinking in mice (13), whereas increasing K_Ca_2 channel function with positive modulators reduced home cage drinking and operant self-administration (11, 17). Importantly, the ability of apamin to inhibit K_Ca_2 channel function was completely lost in NAcC MSNs from ethanol dependent mice, but not in rats that had access to 7-weeks of operant self-administration of moderate amounts of ethanol (11). Together, these studies identified *KCNN3* as a potential regulator of ethanol consumption and revealed NAcC neuroadaptations in K_Ca_2 channel function that may drive excessive drinking in preclinical rodent models of chronic ethanol exposure.

There are two polymorphic CAG repeats in the N-terminus of human *KCNN3* encoded by exon 1 (18). Higher numbers of CAG repeats reduced K_Ca_2 channel function in transfected HEK293 cells (19), and this polymorphism has been associated with neuropsychiatric conditions, such as schizophrenia and anorexia nervosa (18, 20–23). While this polymorphism did not confer risk for developing the disease, longer CAG repeat length associated with higher cognitive performance in individuals with schizophrenia (19). This finding is consistent with a known role for K_Ca_2.3 channel regulation of cognitive function and plasticity of intrinsic excitability that is an important mechanism for new learning to occur (19, 24, 25). Because the CAG trinucleotide repeat is conserved in non-human primates (19, 24–26) and non-human primates exhibit a range of ethanol drinking that mimics human consumption (27–29), the present study explored the relationship between CAG repeat length and ethanol drinking in rhesus macaques with low-and high-drinking phenotypes.

DNA methylation (DNAm) is an epigenetic mark that can contribute to modulation of gene expression by modifying the compaction status of the chromatin. Alterations in DNA methylation are reported in heavy drinking monkeys and rodents (30–32), and a recent study demonstrated that knockdown of DNA methyltransferases reduced *Kcnn3* expression and increased intrinsic excitability of cultured cortical neurons (33). Thus, we measured DNAm levels at a differentially methylated region (DMR) in exon 1 that coincides with a cross-species regulatory region within the *KCNN3* promoter. In humans, the *KCNN3* gene encodes four known transcript variants (TVs) by making use of alternative first exons and alternative splicing. Similar to the CAG trinucleotide repeat, alternative *KCNN3* TVs influence function of K_Ca_2 channels. *KCNN3* variant *SK3_1B* encodes a truncated channel that functions as a dominant-negative to suppress endogenous K_Ca_2 channel currents (34), whereas variant *hSK3_ex4* encodes a protein with an additional 15 amino acid insertion within the S5-PHelix loop that renders the channel insensitive to apamin block (35). We also determined if ethanol-induced changes in DNAm of exon 1 altered expression of *KCNN3* TVs in the NAcC of drinking mice and monkeys. Here, we report a complex cross-species relationship between NAcC *KCNN3* and excessive ethanol drinking that ultimately leads to reduced K_Ca_2.3 channel protein expression in mice and monkeys.

## Materials and methods

### Ethanol self-administration in rhesus macaques

Male and female rhesus macaques (*n* = 66, *Macaca mulatta*) from seven different cohorts (cohorts 4, 5, 6a, 6b, 7a, 7b, and 10) were included in this study (Supplemental Table 1). All of the rhesus macaques were born and reared at the Oregon National Primate Research Center (ONPRC) with their mothers until 2-3 years of age. All subjects were initially selected to minimize relatedness; the average kinship coefficient of all subjects was 0.004. The control subjects were selected by matching age and weight, and then kinship. Controls were housed in the same rooms as the experimental subjects and underwent the same training for awaken blood draws, medical check-ups, and MRI imaging. Controls experienced the same diet, timing and order of experimental phases and had equal experience with the research technicians. However, control monkeys did not have access to ethanol. Instead, controls were yoked to a future ethanol monkey based on weight, and received a quantity of 10% maltose-dextrin solution matched in calories to the previous day’s intake of their yoked ethanol monkey. Monkeys were individually housed and ethanol self-administration was induced using schedule-induced polydipsia, as previously described (29). For all cohorts, monkeys had open access to 4% ethanol and water (ethanol subjects) or water only (control subjects) for 22 h/day for 12 months (see (28) for further details on these seven cohorts). It should be noted that the monkeys in cohort 10 had two cycles of forced abstinence and open access that followed the initial 12 months of open access. Because of the repeated abstinence, samples from monkeys in cohort 10 were limited to *KCNN3* trinucleotide repeat analysis using blood collected prior to ethanol self-administration. While the technicians were not blind to the treatment condition (ethanol or control), the intake data were collected and recorded in an automated fashion by computer and analyzed by individuals who did not interact with or know the drinking status of the monkeys. All of the animal procedures used in this study were approved by the ONPRC IACUC and were performed in accordance with the NIH and the National Resource Council’s *Guide for the Care and Use of Laboratory Animals*.

### Monkey Drinking Phenotypes

The 49 monkeys with access to ethanol were classified into three different age categories based on their age of first ethanol access and four different drinking categories based on previously described criteria (27). There were 15 adolescent (4-5 years of age, ~15-18 human year equivalence; *n* = 6 females), 24 young adult (5-6 years of age, ~18-24 human year equivalence; *n* = 5 females), and 10 mature adult (7-11 years of age, ~25-40 human year equivalence; *n* = 0 females) monkeys. Monkeys were classified as very heavy drinking (VHD) if their average daily ethanol intake was >3 g/kg (~12 drink equivalent in humans) and they consumed >4 g/kg ethanol on ≥ 10% of their open access drinking days. Heavy drinking (HD) monkeys were defined as those that consumed >3 g/kg ethanol on ≥ 20% of their open access drinking days. Binge drinking (BD) monkeys were defined as those that consumed >2 g/kg ethanol on ≥ 55% of their open access drinking days. Low drinking (LD) monkeys were defined as those that did not reach these set thresholds for daily ethanol intake. Classification for cohort 10 was based on their drinking patterns after 12 months of open access. We have previously confirmed that the ethanol drinking behaviors did not reflect general differences in thirst (36).

### Ethanol dependence and two-bottle choice drinking in C57BL/6J mice

Sixty adult male C57BL/6J mice were purchased from Jackson Laboratory (Bar Harbor, ME) at ~7 weeks of age. Mice were individually housed in a temperature and humidity controlled environment and kept on a 12 h light/dark cycle. Food and water were available *ad libitum* during all procedures. The Medical University of South Carolina Institutional Animal Care and Use Committee approved all procedures in accordance with NIH guidelines for the humane care and use of laboratory animals. To establish baseline drinking, half of the mice consumed ethanol in their home cage using a 2-bottle choice (15% ethanol (v/v) vs. water) long-access (22 h) protocol for five weeks. Control mice consumed water in their home cage during this phase. Half of the control and ethanol drinking mice then underwent 4 repeated weekly cycles of chronic intermittent ethanol (CIE) exposure in vapor inhalation chambers, alternated with weekly home cage drinking sessions [2 (water vs ethanol drinking) x 2 (air vs ethanol vapor inhalation) experimental design; *n* = 15 mice/group] with 72 h in between CIE exposure and access to ethanol drinking bottles in their home cages. Ethanol vapor exposure was delivered in Plexiglas inhalation chambers as previously described (13). CIE treatment consists of sixteen hours of vapor exposure followed by eight h of withdrawal. Chamber ethanol concentrations were monitored daily and air flow was adjusted to maintain ethanol concentrations within a range that yields stable blood ethanol levels (175–225 mg/dl) throughout exposure. Prior to entry into the ethanol chambers, EtOH mice were administered ethanol (1.6 g/kg; 8% w/v; i.p.; 20 ml/kg dose volume) and the alcohol dehydrogenase inhibitor pyrazole (1 mmol/kg). Control mice were handled similarly, but received injections of saline and pyrazole. Seventy-two h following the last vapor chamber exposure, mice were given limited access to ethanol or water for 2-3 days prior to sacrifice and tissue collection.

### Genomic DNA and total RNA isolation

After the 12 month open access period, a detailed necropsy protocol was used to systematically collect tissues from all macaques (37). Monkeys were sedated with ketamine (15 mg/kg), and then the animals were brought into a surgical plane of anesthesia with intravenous administration of sodium pentobarbital (30–50 mg/kg, i.v.). Following extraction, the entire brain was blocked or sectioned into slices that were fresh frozen and stored at −80° C. The NAcC was later dissected from frozen brain slices. The typical postmortem interval for brain extractions was < 5 minutes. Mouse brains were rapidly removed and placed in ice-cold saline before blocking 1-mm thick sections using a mouse brain block (ASI Instruments, Inc., Warren, MI, USA). A 1-mm tissue punch (Ted Pella, Inc., Redding, CA, USA) was used to extract the NAcC. Genomic DNA and RNA were extracted from male and female monkey and male mouse NAcC samples using the All Prep DNA/RNA/miRNA Universal kit (QIAGEN Sciences Inc, Germantown, MD) following the manufacturer’s recommendations. Blood samples drawn from macaques prior to ethanol self-administration were used for CAG repeat analysis. Briefly, blood was collected in EDTA tubes and DNA was isolated using QIAamp DNA mini kit following manufacturer’s instructions (QIAGEN Sciences Inc).

### Trinucleotide repeat analysis

Blood DNA was used to analyze the number of CAG repeats within the second *KCNN3* CAG repeat region as previously described (38). The primers (Forward: CAGCAGCCCCTGGGACCCTCG, Reverse: GGAGTTGGGCGAGCTGAGACAG) generated amplicons ranging between 112 bp and 178 bp, depending on the number of CAG repeats present. The PCR products were applied to an ABI3730 XL DNA Analyzer for separation and detection, incorporating a 600LIZ size standard with the PCR products at a 1:1 ratio (Applied Biosystems Inc.). The output files were visualized and product sizes were determined using Gene Mapper 4.0 software (Applied Biosystems Inc.). The sequence content of the amplification products was confirmed by PCR amplifying DNA from individuals with homozygous genotypes, and by direct DNA sequencing of PCR products. We found that the first exon 1 CAG repeat was not variable in rhesus macaques; thus, these studies focused on the second exon 1 CAG repeat length.

### Bisulfite amplicon sequencing

Bisulfite amplicon sequencing was used to measure the DNAm rates of a DMR within the *KCNN3* promoter region using NAcC tissue from macaques and mice. Correction for bisulfite-converted PCR bias was carried out as described by Moskalev et al. (39). Methylated and unmethylated human gDNAs (Zymo Research, Irvine, CA) were bisulfite-converted using the EZ DNA Methylation-Gold kit (Zymo Research, Irvine, CA), according to the manufacturer’s instructions. The bisulfite converted DNAs were combined to create a series of methylation rate standards ranging from 100% to 0% methylated. gDNA (500ng) from rhesus and mouse NAcC was bisulfite converted following the same protocol. Primers were designed to amplify a 646bp region of the *KCNN3* within exon 1A and intron 1 in human, rhesus macaque and mice. Because of the length of the region, two sets of primers were designed to cover the whole region (Supplemental Table 2). Amplification was carried out in the C1000 Thermal Cycler (Bio-Rad, Hercules, CA) using 20 ng of bisulfite-treated DNA per PCR reaction. Amplification was carried out as follows: Phase 1: 10 cycles of 94°C for 30sec., 68° C for 1min., with a touchdown decrease of 1°C per cycle. Phase 2: 28 cycles of 94°C for 30sec. and 58°C for 45sec. Libraries were prepared using the NETflex DNA Sequencing Kit (BIOO Scientific, Austin, TX) according to manufacturer’s instructions. The libraries were evaluated using an Agilent 2100 Bioanalyzer (Agilent Technologies, Palo Alto, CA) and were normalized to 2nM with 10 mM Tris-HCl. The libraries were then pooled and sequenced on a MiSeq (Illumina, Inc. San Diego, CA) by the Molecular & Cellular Biology Core, (ONPRC, Beaverton, OR) to generate 250-base paired-end reads. The reads were trimmed using Trim Galore and aligned to the corresponding reference genome (Rhesus: MacaM (40) and Mouse: GRCm38.p3; https://www.ncbi.nlm.nih.gov/assembly/GCF_000001635.20/) using Bismark (41). M-bias plots were then generated (42), and reads were trimmed further as needed. Bismark alignment data was converted to CpG methylation rate using the Bismark methylation extractor (41) and custom scripts.

To assess the linearity of the PCR amplification we plotted the observed versus expected methylation rates obtained from the methylated DNA standard dilution series (39). Next, a hyperbolic function was applied to correct for PCR-bias obtaining a good linear fit (r_pair-1_^2^=0.94; r_Pair-2_^2^=0.94; r_Pair-3_^2^=0.98; Supplemental Fig. 1).

### RNA isolation and reverse transcription

The NAcC RNA quantity and quality was evaluated on a 2100 Bioanalyzer (Agilent Technologies, Santa Clara, CA). The Fluidigm Reverse Transcription Master Mix (Fluidigm, Inc., San Francisco, CA) was used to reverse-transcribe 100 ng of each RNA sample following the manufacturer’s instructions. Briefly, 1 μl of 100 ng of RNA was combined with 1 μl of Reverse Transcription Master Mix and 3 μl of RNase-free water. The reactions were incubated at 25 C for 2 min, followed by 42 C for 30 min and 85 C for 5 min. Next, 1.25 μl of cDNA was pre-amplified using 1 μl of PreAmp master mix, 0.5 μl of pooled primers (at 500 nM) and 2.25 μl of RNase-free water. The Pre-Amplification conditions were 95 C for 2 min, followed by 10 cycles at 95 C for 15 s and 60 C for 4 min. To remove unincorporated primers, the reaction were mixed with 0.2 μl Exonuclease I reaction buffer, 0.4 μl Exonuclease I (20 Units/ml) and 1.4 μl of RNase-free water. The reactions were incubated at 37 C for 30 min followed by 80 C for 15 min. The reactions were diluted (10x) with 43 μl of TE buffer (TEKnova, Hollister, CA).

### High-throughput real time PCR

qPCR was performed using the BioMark™ HD System and the 96.96 GE Dynamic Arrays (Fluidigm, Inc., San Francisco, CA) in triplicates assays. 5 μL of Fluidigm sample premix consisted of 2.25 μL of 10x diluted pre-amplified cDNA, 0.25 μL of 20x SG loading reagent (Fluidigm), 2.5 μL of Sso Fast Eva Green Mastermix (Bio-Rad). Each 5 μL assay premix consisted of 0.25 μL of 100 μM primers (final concentration 500 nM primers), 2.5 μL 2x Assay loading reagent (Fluidigm) and 2.25 μL of 1x DNA suspension buffer (TEKnova, Hollister, CA). The samples and assays were mixed using the Nanoflex IFC controller (Fluidigm). Thermal qPCR conditions were: 95°C for 60 s, 35 cycles of 95°C for 5 s, and 60°C for 20 s plus melting curve analysis. Data was processed by automatic threshold for each assay, with derivative baseline correction using BioMark Real-Time PCR Analysis Software 3.1.2 (Fluidigm). The quality threshold was set at the default of 0.65.

The primer sequences are described in Supplemental Table 3. Since most of the alternative transcript variants are not annotated in the Rhesus or mouse genome, we used the human annotations to design the primers, then identified the homologous sequence in the rhesus macaque (MacaM) (40) and mouse (GRCm38.p3) genome. The mRNA expression levels were normalized using the phosphoglycerate kinase *(PGK1)* gene. This gene was demonstrated to be a reliable control for brain gene expression (43). We also previously confirmed that different levels of ethanol use did not affect its expression (31). The monkeys 10171 (male) and 10068 (female) and mouse 16C control subjects were used as a reference sample for comparison.

### Western blot analysis

After extraction, tissue samples containing the NAcC extracted from female control and long-term drinking rhesus macaques (*n* = 5/group) were snap frozen, shipped overnight on dry ice, and homogenized in 100 µl of ice-cold homogenization buffer (50 mM Tris-HCl, 50 mM NaCl, 10 mM EGTA, 5 mM EDTA; 2 mM sodium pyrophosphate, 1 mM activated sodium orthovanadate, 0.2 mM AEBSF, 1 µg/ml aprotinin, 1 mM benzamide, 10 µg/ml leupeptin, 10 µg/ml pepstatin, pH 7.5). Samples were probe sonicated for ~ 5 sec and centrifuged at 23,100 × g for 30 min at 4°C. The resulting supernatant was removed and the pellet was resuspended in 2% lauryl dodecyl sulfate (LDS) and probe sonicated for ~ 5 sec. An aliquot was taken for determination of protein concentration by the bicinchoninic acid assay (Pierce Biotechnology, Inc., Rockford, IL). Samples were diluted with NuPAGE 4X LDS sample loading buffer (Invitrogen Corp., Carlsbad, CA; pH 8.5) containing 50 mM dithiothreitol, and samples were denatured for 10 min at 70°C. We first performed a series of western blots using different titrations of sample and antibody to establish the linear range for K_Ca_2.3 (Alomone Labs, Jerusalem, Israel; Catalog #: APC-025; epitope AA 2-21 of human K_Ca_2.3) in primate tissue samples. Specificity of this antibody for K_Ca_2.3 channels has been confirmed in conditional K_Ca_2.3 knockout mice (44). Ten µg of each experimental sample was separated using the Bis-Tris (375 mM resolving buffer and 125 mM stacking buffer, pH 6.4; 7.5% acrylamide) discontinuous buffer system with MOPS electrophoresis buffer (50 mM MOPS, 50 mM Tris, 0.1% SDS, 1 mM EDTA, pH 7.7). Protein was then transferred to Immobilon-P PVDF membranes (Millipore, Bedford, MA) using a semi-dry transfer apparatus (Bio-Rad Laboratories, Hercules, CA). Blots were then washed with phosphate-buffered saline containing 0.1% Tween 20 (PBST) and blocked with PBST containing 5% nonfat dried milk (NFDM) for 1 h at room temperature with agitation. The membranes were then incubated overnight at 4°C with primary antibody diluted 1:4000 in PBST containing 0.5% NFDM and washed in PBST prior to 1 h incubation at room temperature with horseradish peroxidase conjugated secondary antibody diluted 1:2000 in PBST. Membranes received a final wash in PBST and the antigen-antibody complex was detected by enhanced chemiluminescence using a ChemiDoc MP Imaging system (Bio-Rad Laboratories, Hercules, CA). The bands in the experimental samples were background subtracted and quantified by mean optical density using computer-assisted densitometry with Image Lab software (v4.0.1, Bio-Rad Laboratories) with the experimenter blind to the ethanol drinking groups. Because controls (e.g., actin, GAPDH) used to normalize protein loading in western blot experiments can cause quantitation errors (45–47), we used normalized to a total protein following our published methods in monkey tissue (48).

### Statistical Analysis

Data from heavy and very heavy drinking monkeys were combined due to small sample sizes in the transcript analysis and bisulfite sequencing studies. All statistical analyses were carried out using IBM SPSS Statistics (Armonk, NY) except where noted, with values α<0.05. The Shapiro-Wilk test (appropriate for small sample sizes) was used to assess the normality of the average methylation rate, mRNA expression rate, and K_Ca_2.3 protein expression level per comparison group. All variables analyzed followed a normal distribution. Welch’s one-way ANOVA was used to compare the difference in average methylation between controls and ethanol drinkers with Games-Howell post-hoc tests. One-way ANOVA was used to compare mRNA relative expression levels between groups. Prior to applying one-way ANOVA, Levene’s test was used to test homogeneous variance assumption for parametric methods. When heterogeneous variance was detected, we used the nonparametric Kruskal-Wallis test. Bonferroni or Tukey correction for the multiple comparisons were used to correct the overall type I error rate. Two-tailed independent t-test was used to compare the difference in average methylation rate between controls and dependent mice. Based on the Levene’s test for homogeneous variance, we used the appropriate p-value (homogeneous or heterogeneous variance). The allele frequency distribution of CAG trinucleotide repeats between controls and drinking monkeys was compared using the Kruskal-Wallis test (GraphPad Prism software, version 7.04, La Jolla, CA). Normalized western blot data were analyzed by a two-tailed t-test in Prism. Ethanol drinking data in mice were analyzed by a repeated-measures mixed linear model with a Tukey post-hoc test (SAS Institute, Cary, NC, USA).

## Results

### Description of the animal ethanol drinking patterns

Sixty-six unrelated, male and female rhesus macaques were used in this study. In addition to the 16 control monkeys, there were 16 LD, 9 BD, 9 HD, and 16 VHD monkeys. The average daily ethanol intake (range: 0.47 to 5.15 g/kg) across the 12 months of open access for each of the drinking monkeys along with their age at drinking onset and % of drinking days over 3 g/kg are shown in Fig. 1a. While the adolescent and young adult monkeys displayed a wide range of intake, only one of the mature adult monkeys used in this study met criteria for HD or VHD. More detailed analyses of their drinking patterns have been reported previously (27, 28).

**Fig. 1.**
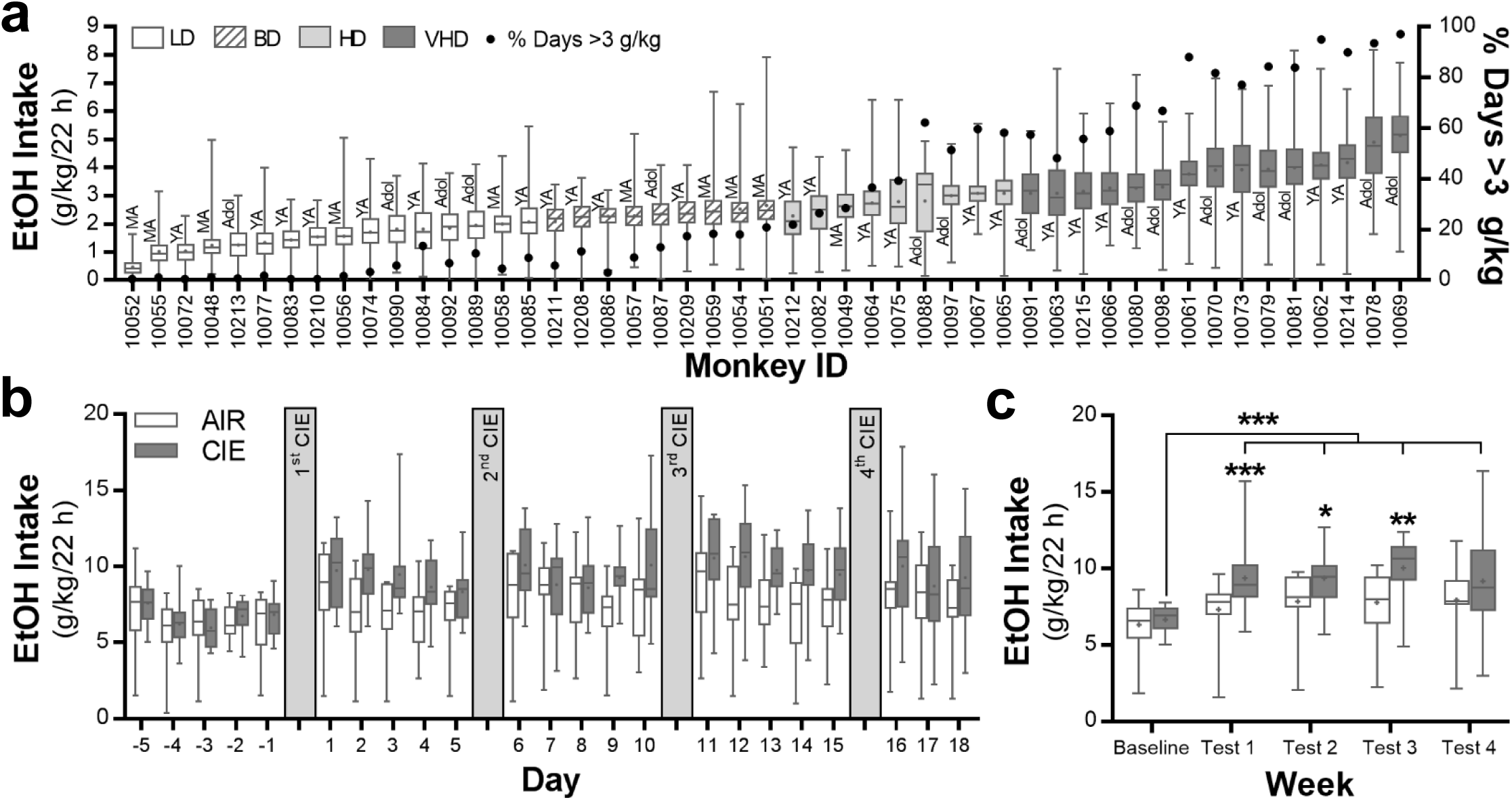
Ethanol consumption for the monkeys and mice using in the current study. **a** Average daily ethanol (4% v/v) intake in male and female rhesus macaques and their drinking category during 12 months of open access. Also shown is their age at the time of first ethanol exposure and the % days where they consumed >3 g/kg/22 h. Adol, adolescent; BD, binge drinking; HD, heavy drinking; LD, low drinking; MA, mature adult; VHD, very heavy drinking; YA, young adult. **b** Daily ethanol (15% v/v) intake values in male C57BL/6J mice prior to and after each of four cycles of chronic intermittent ethanol (CIE) exposure in vapor inhalation chambers (n = 15 mice/group). **c** Average weekly ethanol intake in mice during the week prior to and after each cycle of CIE exposure (Tukey post hoc, \*\*\**p* = 0.0074 vs test 1 air group; \**p* = 0.0468 vs test 2 air group; \*\**p* = 0.0034 vs test 3 air group; \*\*\**p* = <0.0001 tests 1-4 vs baseline CIE group).

To more closely match the monkey drinking paradigm, the standard two-bottle choice, limited-access mouse model of dependence-induced escalation of drinking was modified to allow mice to have access to ethanol (15% v/v) for 22 h/day. Ethanol drinking behavior prior to and following each weekly exposure to CIE is shown in Fig. 1b,c (*n* = 15 mice/group). Consistent with studies using limited access to ethanol (13, 49, 50), mice exposed to CIE significantly increased their voluntary ethanol intake (interaction: F(4,109) = 2.47, *p* = 0.0491). Post hoc analysis indicated that the two treatment groups were not different at baseline (p = 0.6685), but differed during weekly test drinking sessions 1 (*p* = 0.0074), 2 (*p* = 0.0468), and 3 (*p* = 0.0034). In the ethanol dependent mice, drinking levels in all four test sessions were significantly higher than their intake during baseline (*p* = <0.0001).

### *KCNN3* methylation analysis

By comparing the DNA sequence of the *KCNN3* gene and promoter across human, rhesus macaque, and mouse, we identified ten regions with potential conserved regulatory region, based on sharing over 95% sequence homology and ≥ 75% CpG identity. We then analyzed the DNAm patterns in these ten candidate regions between ethanol-naïve, LD, and HD/VHD rhesus macaques. Because of the small sample size, BD were not included in this analysis, while HD and VHD were combined based on their similar drinking behavior. Among the different regions, only a region of 646 bp overlapping with exon 1 and intron 1 of *KCNN3* (MR-ex1; Fig. 2a) showed significant DNAm differences between groups (Fig. 2b). The MR-ex1 region contained 24 CpGs in rhesus macaques that were 96% conserved in humans (Supplemental Fig. 2). While the overall CpG conservation was lower in the mouse as compared to both human and rhesus macaques (15 CpGs, 75%), the high sequence and CpG similarity of this region across species suggests functional relevance and underscores the potential translational value of the DNAm signal identified in this study. Overall, the CpGs within MR-ex1 showed generally low DNAm in controls, with methylation levels ranging from 7-24% in males (Fig. 2b) and 3-28% in female macaques (Fig. 2c). In males, LD monkeys showed similar DNAm levels as controls; however, HD/VHD monkeys had increased methylation rates as compared to both controls and LD monkeys (Fig. 2b). In particular, nine CpGs had significantly higher methylation rates in male HD/VHD monkeys compared with control monkeys. In females, LD subjects could not be included in the analysis due to the small sample size (only 3 subjects). Nonetheless, and similar to heavy ethanol-drinking males, there were four CpGs with significantly higher methylation rates in the MR-ex1 region of VHD female macaques as compared to ethanol-naïve females (Fig 2c).

**Fig. 2.**
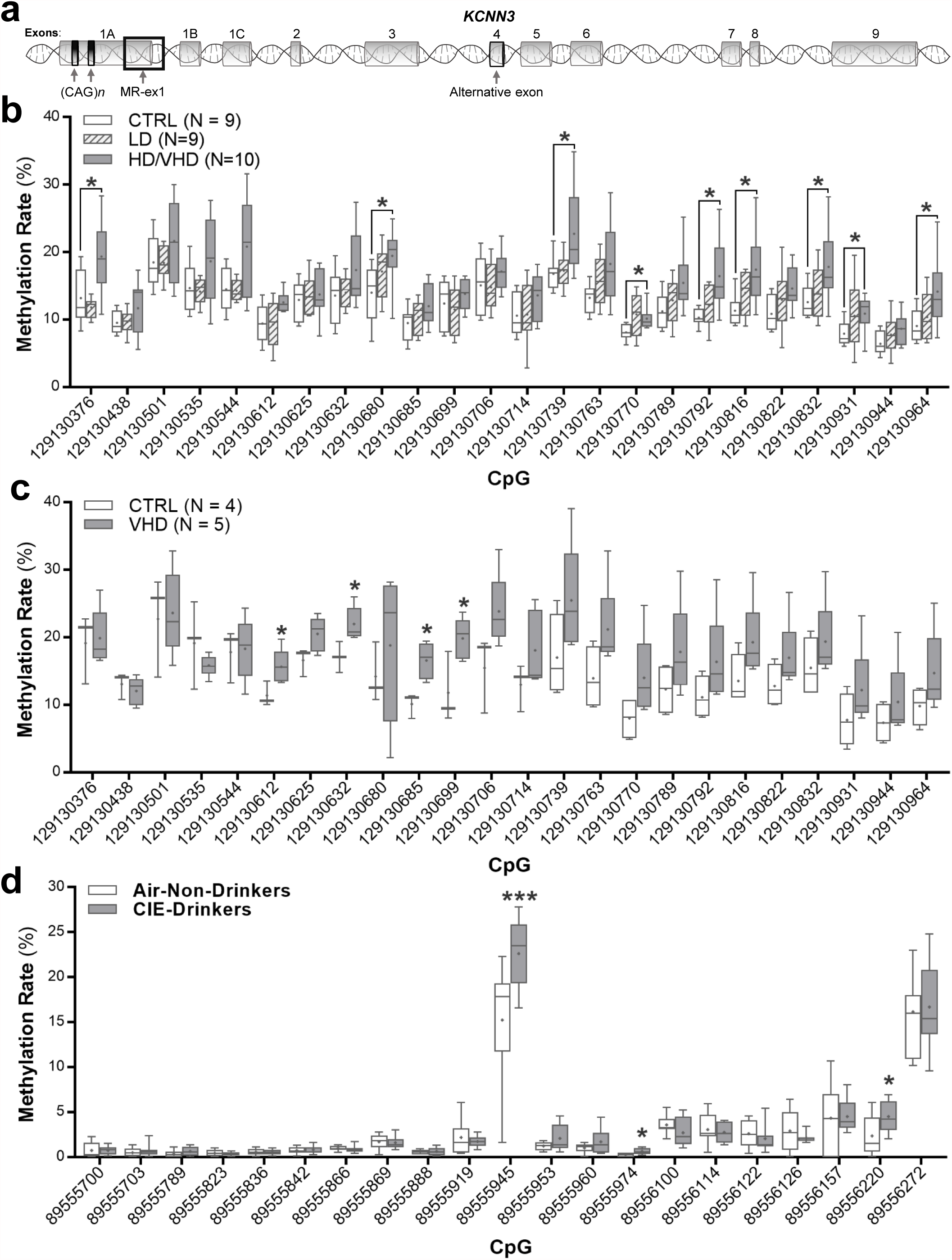
*KCNN3* methylation levels within MR-ex1-200 of ethanol drinking monkeys and dependent mice. The average methylation rates of individual CpGs included in the methylation region under study are shown. **a** Exon organization of the *KCNN3* locus showing the location of exons and the dual CAG trinucleotide repeat arrays and methylated region in exon 1 (MR-ex1). **b** In rhesus macaque, the following CpGs showed elevated rates of methylation in heavy/very heavy drinking macaques vs controls: CpG_129130376_: F(2, 11.846) = 7.9, \**p* = 0.04; CpG_129130680_: F(2, 15.955) = 3.661, \**p* = 0.036; CpG_129130739_: F(2, 15.468) = 3.817, \**p* = 0.04; CpG_129130770_: F(2, 13.57) = 5.047, \**p* = 0.041; CpG_129130792_: F(2, 13.945) = 7.047, \**p* = 0.01; CpG_129130816_: F(2, 15.63) = 5.836, \**p* = 0.015; CpG_129130832_: F(2, 15.473) = 3.95, \**p* = 0.033; CpG_129130931_: F(2, 14.393) = 4.016, \**p* = 0.038; CpG_129130964_: F(2, 15.708) = 3.985, \**p* = 0.031. **c** In female macaques, elevated rates of methylation were observed at the following CpGs in very heavy drinking macaques vs controls: CpG_129130612_: F(1, 7) = 6.122, \**p* = 0.048; CpG_129130632_: F(1, 7) = 7.64, \**p* = 0.033; CpG_129130685_: F(1, 7) = 13.370, \**p* = 0.011; CpG_129130699_: F(1, 7) = 7.799, \**p* = 0.031. **d** In mouse accumbens, the following CpGs showed elevated rates of methylation between non-drinking mice and drinking dependent mice: Independent t-tests: CpG_89555945,_ *t*(16)= −2.946, \*\*\**p* = 0.0009; CpG_89555974,_ *t*(17) = −2.860, \**p* = 0.011; CpG_89556220,_ *t*(17) = −2.206, \**p* = 0.042.

Comparison of the MR-ex1 region to human ENCODE data (51) on 25 chromatin states for seven different brain areas predicted that this region coincides with promoter function (Supplemental Fig. 3). Furthermore, and in agreement with a potential role of this region as a promoter (52), several transcription factors relevant to neuronal regulation and known to have a role in mediating the effects of ethanol on gene regulation are predicted to bind to it, including GR (glucocorticoid receptor; (53)), ER-α (estrogen receptor; (54)), CREM (cAMP responsive element modulator; (55)), CREB (cAMP responsive element binding protein; (56)), Sp1 (57), GATA-3 (58), NeuroD1 (59), C/EBP (CCAAT-enhancer-binding proteins; (60)) and AP-2α (61) (Supplemental Fig. 4, TRANSFAC (62)). In addition to the fact that this methylation region is located in exon 1 (605 bp downstream of the transcription start site of exon 1A), it is upstream of exons 1B and 1C (~28kb and ~2kb; respectively), and could act as a regulatory region contributing to differential expression of *KCNN3* TVs.

We next investigated the methylation profile of the MR-ex1 region in mice that were drinking ethanol for 22 h in the CIE dependence model. Similar to ethanol-naïve rhesus macaques, this region was hypomethylated in air-exposed control mice. *Kcnn3* methylation levels ranged from 0.1% to 17%, with average methylation rates for 19 out of 21 CpGs below 5% in the controls. Interestingly, three CpGs showed significantly higher methylation rates in CIE-exposed drinking mice as compared to rates in the controls (Fig. 2d). These CpGs are conserved with rhesus macaques and humans (Supplemental Fig. 2) and two were located in the binding sites for AP-2α (Supplemental Fig. 4). Importantly, one CpG with predicted AP-2α, CREB, and GATA-3 binding sites was significantly differentially methylated in both female HD/VHD macaques and CIE-exposed mice.

### Expression of *KCNN3* transcript variants differs with ethanol intake levels

We next evaluated the potential relevance of MR-ex1 hypermethylation in regulating *KCNN3* mRNA expression. The mouse (GRCm38.p3) and rhesus macaque (MacaM) genomes are not annotated with as much detail as the human. Thus, in order to investigate the effects of ethanol drinking on the expression of the different *KCNN3* TVs, we designed primers to amplify two exons common to all reported TVs (*SK3_ex7/8*), as well as primers to specifically amplify TV *SK3_ex4*, *SK3_ex1B*, and *SK3_ex1C* in human. Next, each amplicon’s orthologous sequence was identified in mouse and rhesus macaque, and species-specific primers for the different TVs were designed. It should be noted that TV *SK3_ex1C* was not detected in the NAcC, in agreement with previous studies indicating this TV is not expressed in the brain (63).

The expression of the exons common to all TVs (i.e., *SK3_ex7/8*) showed no differences among the different ethanol-drinking male monkey groups (one-way ANOVA: *F*(2, 23) = 2.721, *p* = 0.0870, *n* = 8-9/group; Fig. 3a). However, male monkeys with a HD/VHD phenotype showed a significant increase in expression of TV *SK3_ex1B* (one-way ANOVA: *F*(2, 22) = 3.547, *p* = 0.0462, *n* = 7-10/group) and *SK3_ex4* (one-way ANOVA: *F*(2, 22) = 9.89, *p* = 0.0009, *n* = 8-9/group) that encode dominant-negative and apamin-insensitive isoforms of K_Ca_2.3 channels, respectively (Fig. 3b,c; Supplemental Table 4). In addition, TV *SK3_ex4* was significantly increased in the accumbens of LD male monkeys (Fig. 3c). Since the male monkeys differed in their age of drinking onset, we wanted to determine if age is an important factor in ethanol regulation of *KCNN3* transcript variant levels. Because of the lack of mature adult control monkeys, we could only include samples from adolescent and young adult male monkeys. In controls, expression of the TVs was not significantly different (*SK3_ex7/8*: two-tailed *t*-test; *t*(2) = 1.802, *p* = 0.2134; *SK3_ex1B*: two-tailed *t*-test; *t*(2) = 0.1494, *p* = 0.8950; *SK3_ex4*: two-tailed *t*-test; *t*(2) = 0.6137, *p* = 0.6019); thus, samples from these two age groups were pooled for further analysis. When collapsed across age, 12 months of open access to ethanol regardless of drinking phenotype significantly increased *KCNN3* TV *SK3_ex4* expression (one-way ANOVA: *F*(2, 17) = 10.88, *p* = 0.0009; Fig. 3f), but not expression of the two other TVs (*SK3_ex7/8*: one-way ANOVA: *F*(2, 18) = 1.798, *p* = 0.1941; *SK3_ex1B*: one-way ANOVA: *F*(2, 17) = 2.935; *p* = 0.0803; Fig. 3d,e), in adolescent and young adult male monkeys.

**Fig. 3.**
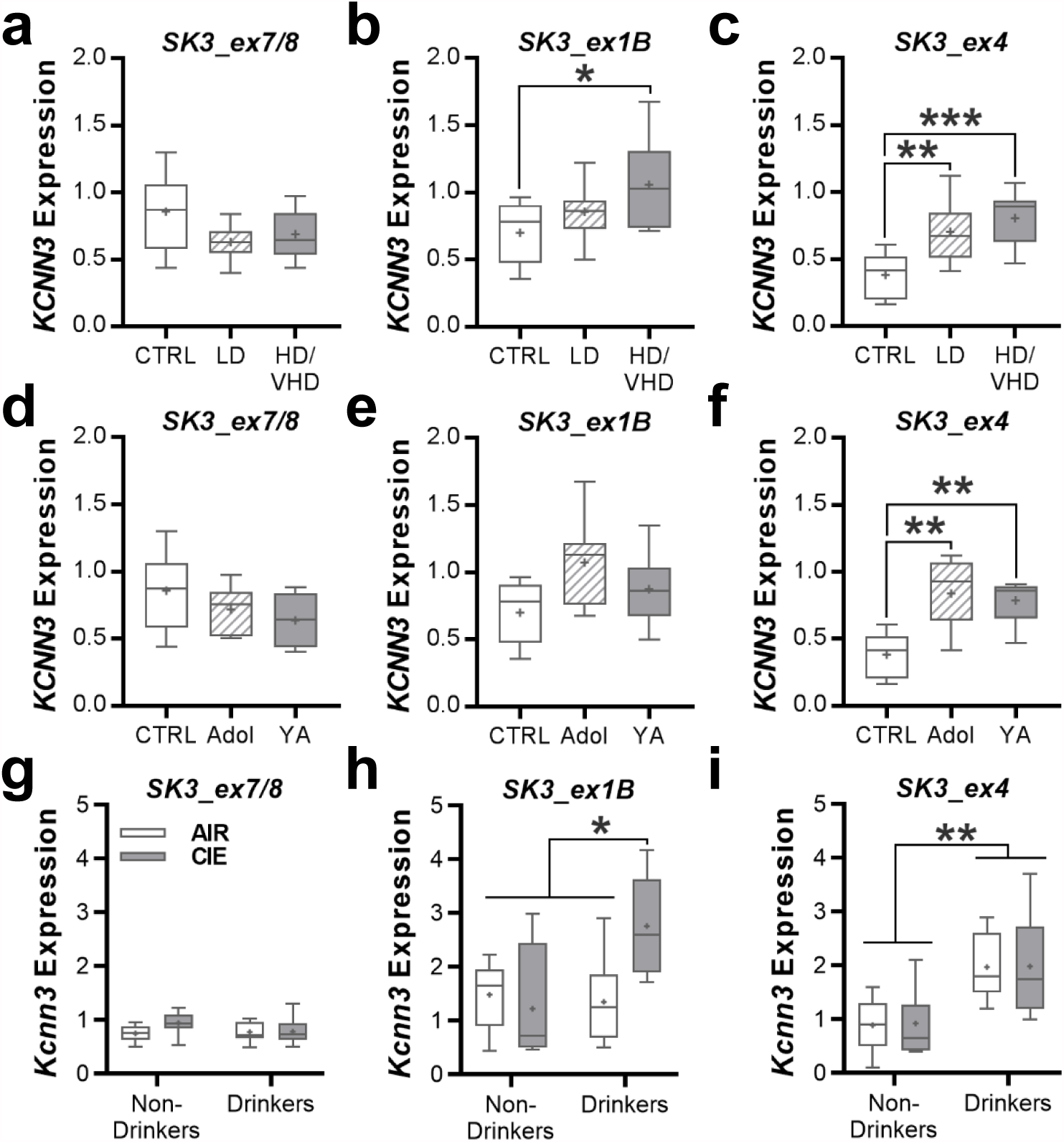
Summary of nucleus accumbens *KCNN3* transcript expression in ethanol drinking rhesus macaques and C57BL/6J mice. **a-c** The relative expression of brain *KCNN3* transcripts (*SK3_ex7/8*, 1 *SK3_ex1B*, and *SK3_ex4*) among the three drinking macaque groups (*SK3_ex1B*: \**p* = 0.0381 vs CTRL; *SK3_ex4*: \*\**p* = 0.0084 vs CTRL, \*\*\**p* = 0.0009 vs CTRL). **d-f** The relative expression of the *KCNN3* transcripts in drinking monkeys collapsed by age at onset of ethanol drinking (*SK3_ex4*: \*\**p* < 0.0015 vs CTRL). **g-i** The relative expression of the *KCNN3* transcript variants in the accumbens of male C57BL/6J mice that were treated with CIE exposure (*SK3_ex1B*: \**p* < 0.026 CIE exposed drinkers vs all remaining groups; *SK3_ex4*: \*\**p* < 0.015 drinkers vs non-drinkers).

Because *KCNN3* is hypermethylated in HD/VHD monkeys and transcriptionally regulated by estrogen (64), analysis of *KCNN3* TVs was also performed in control and ethanol-drinking female monkeys. Similar to the male monkeys, *KCNN3* TV *SK3_ex7/8* expression did not differ between the control and VHD female monkeys (two-tailed *t*-test: *t*(8) = 0.036, p = 0.972, n = 4-6/group; Fig. 4a). Female monkeys with a VHD phenotype showed a significant increase in expression of TV *SK3_ex1B* (two-tailed *t*-test: *t*(10) = 2.658, p = 0.024, n = 5-7/group; Fig. 4b) and *SK3_ex4* (two-tailed *t*-test: *t*(9) = 2.477, p = 0.037, n = 5-6/group; Fig. 4c). All females were of similar age (i.e., 4 – 6 years old at the start of induction), and no age effect on gene expression was performed. To determine if hypermethylation of *KCNN3* and the shifts in TV expression affected K_Ca_2.3 channel protein expression, we performed western blot analysis in accumbens from the same female rhesus macaques. Characterization of K_Ca_2.3 channel immunoreactivity in ethanol-naïve monkey accumbens samples revealed a linear dynamic range across twofold dilutions between 1.25 and 40 μg of protein (R^2^ = 0.9963; Fig. 4d,e). Consistent with results from rodent ethanol studies (11, 13, 17), expression of K_Ca_2.3 channel protein was significantly reduced in VHD female monkeys compared with controls (two-tailed *t*-test: *t*(8)= 2.585, p = 0.0324, *n* = 5/group; Fig. 4f,g). Unfortunately, our attempts to detect the different isoforms of K_Ca_2.3 channel, such as the dominant-negative isoform 3 with a predicted molecular weight of 47 kDa, were unsuccessful with commercially available antibodies.

**Fig. 4.**
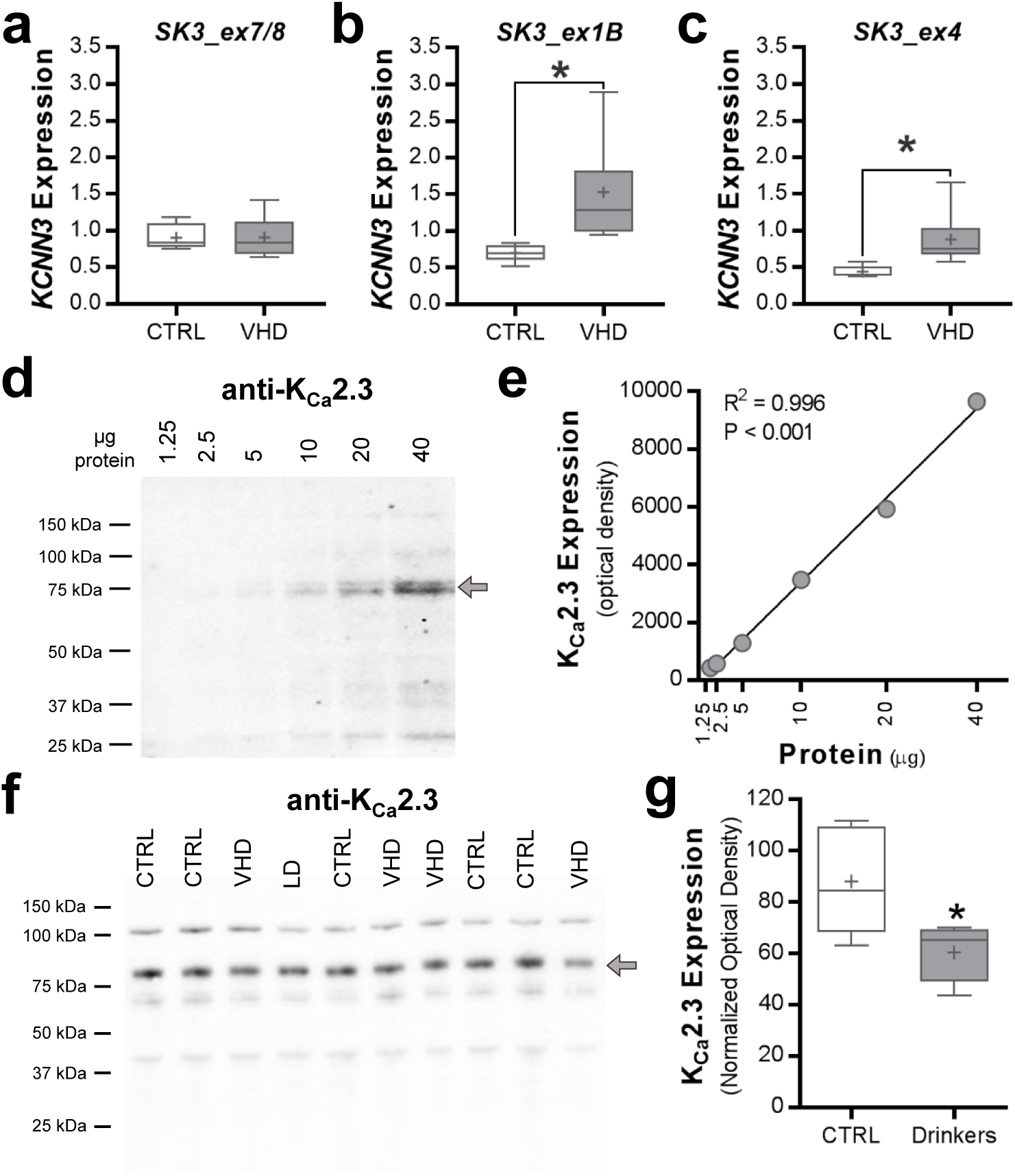
Adaptations in nucleus accumbens *KCNN3* transcript and protein expression in ethanol drinking female rhesus macaques. **a-c** The relative expression of *KCNN3* transcripts (*SK3_ex7/8*, 1 *SK3_ex1B*, and *SK3_ex4*) among control and very heavy drinking macaque groups (*SK3_ex1B*: \**p* = 0.037 vs CTRL; *SK3_ex4*: \**p* = 0.024 vs CTRL). **d** Characterization of anti-K_Ca_2.3 channel western blot in macaque accumbens tissue (protein loading range, 1.25 – 40 µg). **e** Positive correlation between the amount of protein loaded and anti-K_Ca_2.3 channel optical density values. **f,g** The full K_Ca_2.3 channel blot and quantitation of normalized K_Ca_2.3 channel protein expression in controls and drinkers (\**p* = 0.0324 vs CTRL).

Given that chronic ethanol drinking and dependence reduced K_Ca_2 currents and expression in the accumbens of rats and mice (11, 13, 17), we next determined if *Kcnn3* TV expression is altered in ethanol dependent mice. Similar to the monkey data, ethanol drinking and/or dependence did not affect *Kcnn3* expression of *SK3_ex7/8* (two-way ANOVA: main effect: *F*(1, 50) = 3.931, *p* = 0.0529; Fig. 3g). In contrast, dependent mice with access to ethanol in their home cage showed elevated *SK3_ex1B* expression (two-way ANOVA: interaction: *F*(1, 31) = 7.228, *p* = 0.0114; Fig. 3h). Expression of *SK3_ex4* was significantly increased in drinking mice regardless of their history of CIE exposure (two-way ANOVA: *F*(1, 44) = 32.29, p < 0.0001; Fig. 3i).

### *KCNN3* polymorphisms and ethanol consumption levels

In addition to DNAm as a potential regulator of *KCNN3* expression, MR-ex1 contains two polymorphic sites in the promoter region of the *KCNN3* gene that are composed of a variable number of CAG repeats and have been associated with K_Ca_2 channel activity. It has been reported higher numbers of CAG repeats reduces K_Ca_2 channel currents (19). We then investigated the presence of these polymorphic sites in ethanol-drinking rhesus macaques and whether they would influence *KCNN3* regulation. In contrast to the first CAG repeat array that was not polymorphic, the second CAG repeat array was highly polymorphic across rhesus monkeys (Fig. 5a) and ranged from seven to 30 repeats. We found no differences between the frequency distributions for CAG repeats in low, binge, high and very high drinking monkeys (*H*(4) = 2.354, *p* = 0.5023; Fig. 5a). In addition, the sum of CAG repeats of both alleles did not correlate with ethanol intake values (*p* = 0.6946; Fig. 5b), the expression of *KCNN3* variants (*p* ≥ 0.1631; Fig. 5c-e), nor averaged MR-ex1 DNAm rates (*p* = 0.3563; Supplemental Fig. 5).

**Fig. 5.**
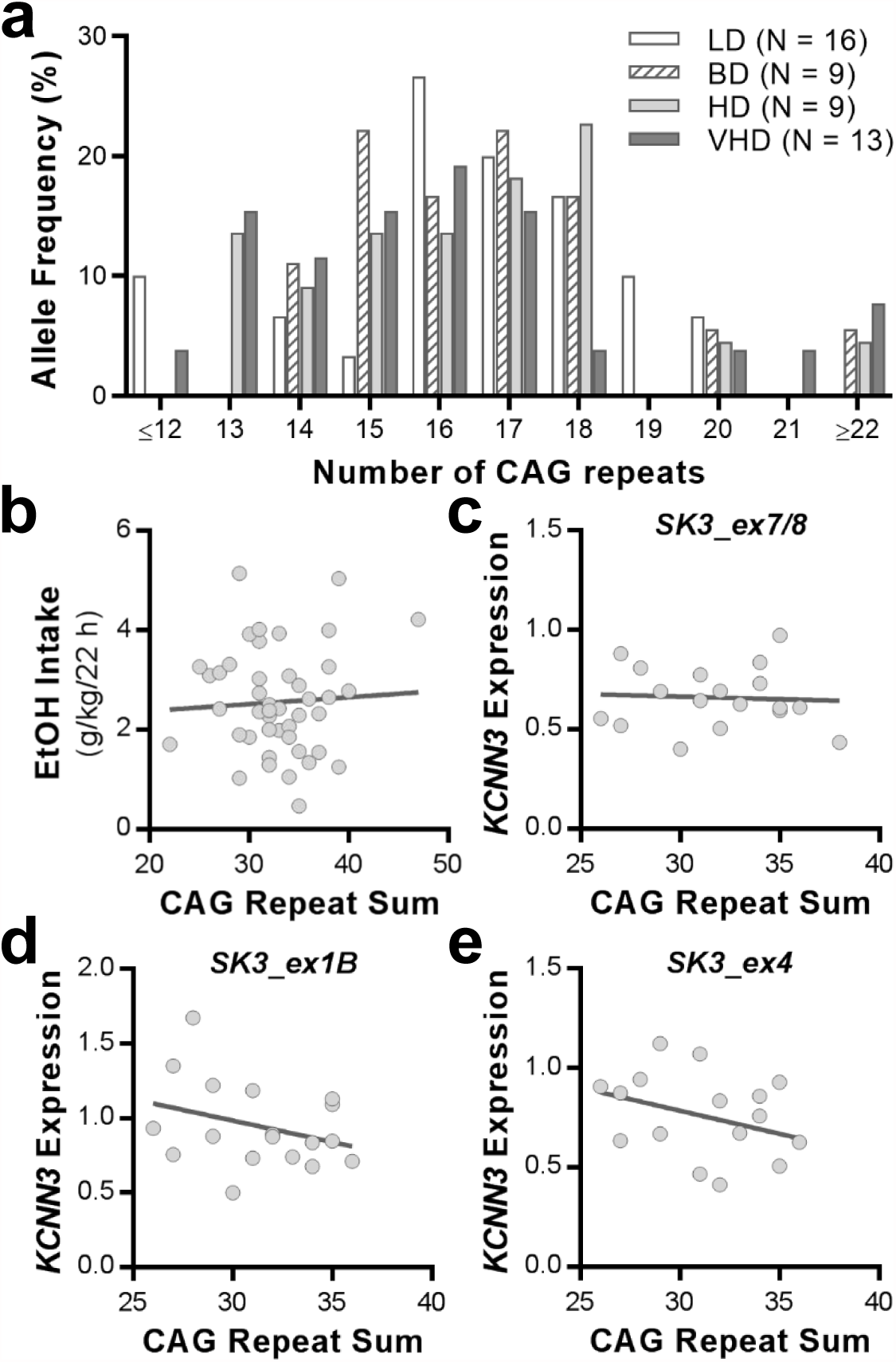
*KCNN3*-CAG*n* allele frequency distribution among male and female rhesus macaques. **a** The frequency distribution of (CAG)*n* alleles is shown for low drinkers (LD), binge drinkers (BD), heavy drinkers (HD), and very heavy drinkers (VHD). **b** Correlation between CAG repeat sum and average ethanol intake. **c-e** Correlations between CAG repeat sum and *KCNN3* transcript expression in long-term drinking rhesus macaques.

## DISCUSSION

The results from these studies provide cross-species evidence for epigenetic regulation of accumbens *KCNN3* expression in heavy drinking macaques and ethanol dependent mice. In both monkeys and mice, ethanol drinking and dependence was associated with hypermethylation of conserved CpGs at a predicted regulatory region in exon 1A of *KCNN3*. In parallel with the hypermethylation, excessive drinking increased expression of a dominant-negative transcript variant of *KCNN3* that is transcribed using an alternative exon downstream of exon 1A. Consistent with chronic ethanol-induced loss of apamin-sensitive currents in accumbens and orbitofrontal neurons (11, 13, 65), ethanol drinking increased expression of the transcript variant that encodes apamin-insensitive K_Ca_2 channels. We also found a reduction in K_Ca_2.3 channel protein in heavy drinking female macaques that is congruent with decreased expression reported in rodent models of chronic ethanol exposure. These results suggest that ethanol-induced regulation of *KCNN3* transcript variants is a conserved mechanism that underlies functional changes in K_Ca_2 channels reported in rodent models.

In the current study, we identified a DMR that maps to an ion channel gene previously implicated in excessive drinking, ethanol-seeking behaviors, and ethanol-induced plasticity of intrinsic excitability (6, 10, 11, 13, 17). Our analysis identified a DMR that spans exon 1 and part of intron 1 of monkey and mouse *KCNN3* in a region containing CpGs that are highly conserved across species. Across male and female monkeys, more than half of the CpGs in this DMR were hypermethylated in HD/VHD, but not in LD or ethanol-naïve monkeys, an effect that is also conserved in ethanol dependent mice that show escalated drinking. Importantly, the DMR in exon 1 of *KCNN3* coincides with a predicted promoter region with transcription binding sites for neural relevant transcription factors. Furthermore, this DMR is upstream of two alternative exons 1 (1B and 1C). All together, these findings suggest that the DMR is strategically located to potentially regulate alternative transcript expression of *KCNN3*. A previous characterization of the promoter region upstream of the exon 1A transcription start site (TSS) identified consensus sequence binding sites for CREB, AP-1, and AP-2 (52). Additional binding sites for these same transcription factors are predicted to bind to MR-ex1, specifically to significantly differentially methylated CpGs. Numerous signaling transduction pathways that are altered by ethanol consumption lead to CREB activation (66), a key mediator in the development of addiction. Upon activation, CREB can exert its influence upon target gene transcription and interact with promoter-bound cofactors. Previous evidence showed that CREB modulates BK channel expression (67), and our results suggest that it may also modulate *KCNN3* expression. AP-1 complexes containing FosB, which accumulates in the NAc after drug intake, modulates promoters of genes relevant to addiction, such as GluA2 and dynorphin (68). The MR-ex1 region also contains binding sites for glucocorticoids, which have been extensively associated with AUD (69, 70). Others have shown that glucocorticoids and stress exert profound effects on intrinsic excitability in neurons through regulation of ion channel activity, including K_Ca_2 channels (71–74). Thus, it is possible that ethanol, by modulating transcription factor levels as well as availability of binding sites by DNAm, modulates *KCNN3* expression. Using a genome-wide approach, we previously reported DMRs associated with the modulation of genes that regulate synaptic plasticity in the NAcC of heavy drinking monkeys (31). In a recent analysis of the NAcC methylome of ethanol-naïve and HD/VHD macaques, we identified the same DMR described in this study (unpublished data). Furthermore, induction of ethanol dependence in C57BL/6J mice increased evoked firing in NAcC MSNs (13) and induced synaptic proteome adaptations in the NAcC (75). Because alterations in neuronal firing underlie synaptic integration and learning processes and may facilitate drug-associated synaptic remodeling (76, 77), our findings suggest that a change in the methylation status of key CpGs is a critical cross-species mechanism that regulates possible coordinated neuronal excitability and synaptic adaptations in heavy drinking animals.

In monkeys and mice, heavy ethanol drinking and dependence increased expression of *KCNN3* transcript variants that encode K_Ca_2.3 channels that reduce surface trafficking and apamin sensitivity. A combination of TV and protein expression data suggest that heavy ethanol consumption results in a decrease in TV *SK3_ex1A* expression while there is an upregulation in TVs *SK3_ex1B* and *SK3_ex4*. While expression of the dominant-negative TV was only increased in heavy drinking monkey, expression of the apamin-insensitive TV was increased by ethanol intake regardless of the drinking phenotype or the age of drinking onset. These results are in agreement with our functional and behavioral data on reduced apamin sensitivity in the nucleus accumbens and orbitofrontal cortex of ethanol dependent mice and self-administering rats (11, 13, 65). We previously reported that apamin microinfusion into the NAcC increased drinking and bath application of apamin reduced K_Ca_2-mediated currents in NAcC MSNs in non-dependent C57BL/6J mice. However, the ability of apamin to influence drinking and K_Ca_2 currents was completely lost when mice were exposed to CIE. Moreover, ethanol intake and evoked firing in NAcC MSNs were increased and there was a reduction in K_Ca_2 channel currents and protein in the ethanol dependent mice. Although it is unknown if heavy ethanol drinking in macaques alters firing properties of NAcC MSNs, the shift in transcript variants and the reduction in total K_Ca_2 channel protein suggests that prolonged ethanol intake will increase intrinsic excitability similar to results from rodent models of ethanol self-administration (11) and dependence (13, 78). In addition to reduced *Kcnn3* gene expression and increased intrinsic excitability, a recent study reported a loss of apamin’s ability to increase evoked firing in cultured cortical neurons treated with DNA methyltransferase inhibitors (33). Thus, these data provide support that the increase in *SK3_ex1B* and *SK3_ex4* TV expression through hypermethylation of *KCNN3* exon 1A is an underlying mechanism driving these functional and behavioral adaptations across species and brain regions.

Previous studies using functional genomics and rodents with divergent drinking phenotypes have identified *Kcnn3* as a candidate signature gene that is associated with binge-like and excessive ethanol drinking (6, 9, 13, 14, 16). In contrast to this mounting evidence and our hypothesis, high numbers of polymorphic trinucleotide repeats encoded by exon 1 of *KCNN3* did not segregate with a heavy drinking phenotype in this population of rhesus macaques. There are a number of possibilities that could explain these negative findings. Although long CAG repeats in K_Ca_2.3 channels reduced apamin-sensitive currents (19), function of this polymorphism was characterized in transfected HEK293 cells and not mammalian central neurons. In the current study, CAG repeat number was measured from blood samples taken after chronic ethanol consumption. While traditionally considered stable, there is some evidence that CAG repeats can vary between types of tissue (i.e, peripheral vs central) and can expand across time to accelerate disease progression (79, 80). Thus, future longitudinal studies are necessary to track CAG repeat number in the NAcC of LD and HD/VHD monkeys.

In summary, these cross-species findings on genetic and epigenetic adaptations in *KCNN3* by excessive alcohol consumption represent a complex mechanism through the use of alternative promoters that likely impact intrinsic excitability of NAcC MSNs, and, ultimately, ethanol-seeking behaviors. We propose a model in which MR-ex1 functions as a regulatory region to modulate the expression of the alternative transcript variants *SK3_ex1B* and *SK3_ex4*. Our findings provide the first evidence that hypermethylation of the MR-ex1 region of *KCNN3* by heavy alcohol drinking is a key cross-species mechanism that may be important for the maintenance of excessive drinking and the development of AUD.

## ACKNOWLEDGEMENTS

The authors would like to acknowledge the support of the Monkey Alcohol Tissue Research Resource (NIH grant AA019431 (KAG)) and Drs. Erich Baker and James Daunais for their assistance with the monkey drinking data and monkey brain samples, respectively. This work was supported by NIH grants AA020930 (PJM), AA013641 (KAG), AA013510 (KAG), AA026092 (RCJ), and AA020928 (BMF).

---

**Supplemental Table 1.**
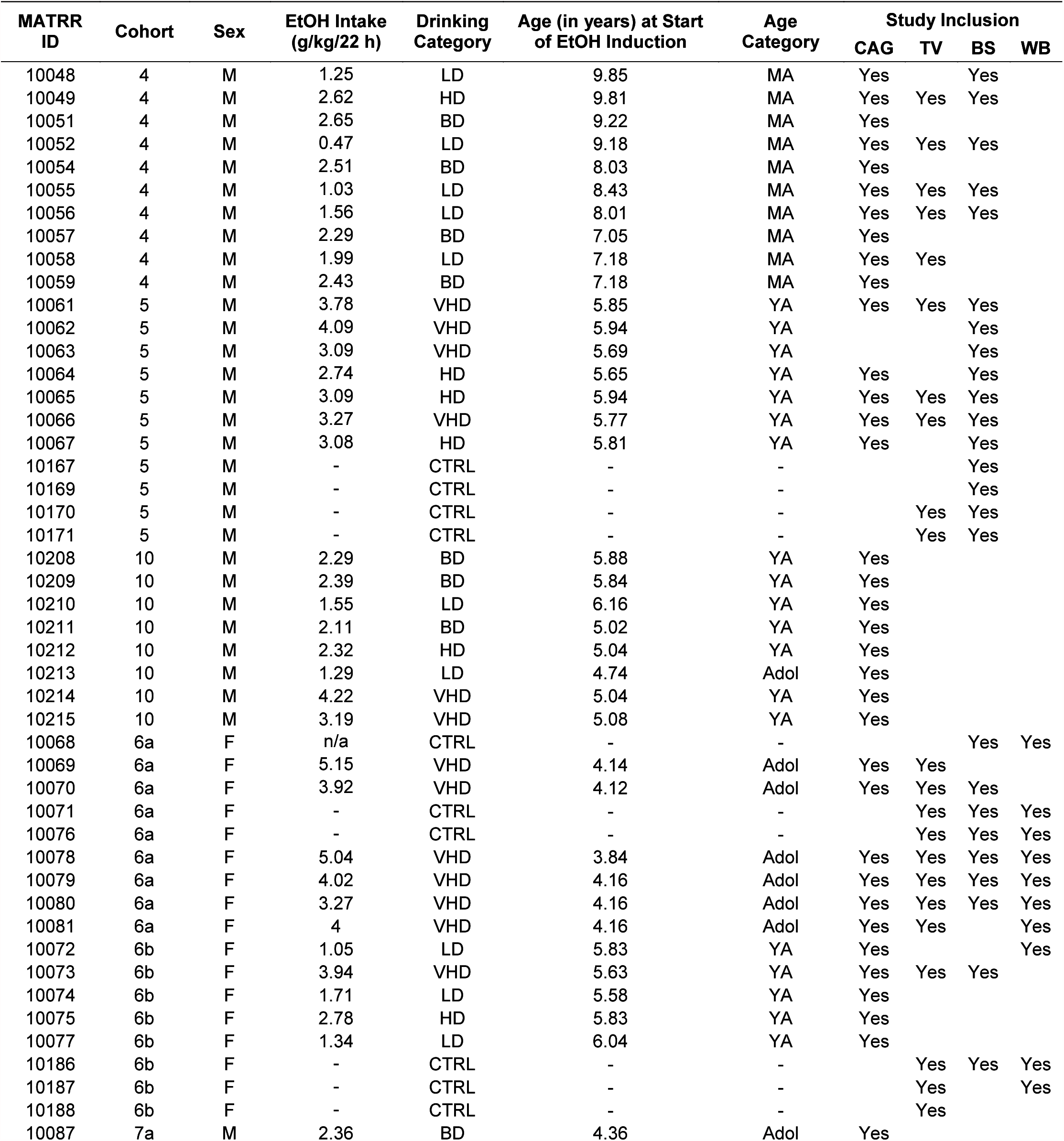
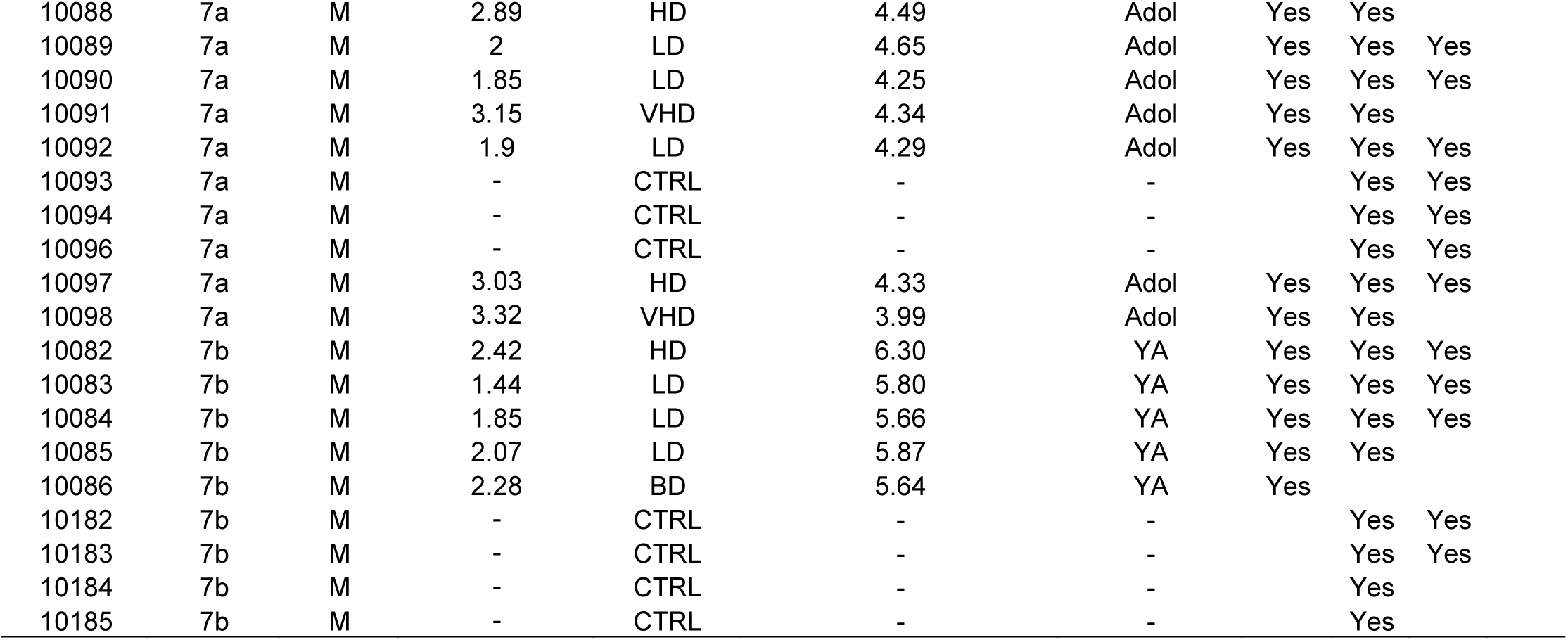
Descriptions of the male and female rhesus macaques used in these studies. Adol, adolescent; BD, binge drinking; BS, bisulfite sequencing analysis; CAG, trinucleotide repeat genotyping analysis; HD, heavy drinking; LD, low drinking; MA, mature adult; TV, *KCNN3* transcript variant analysis; VHD, very heavy drinking; WB, K_Ca_2.3 channel western blotting analysis; YA, young adult.

| MATRR ID | Cohort | Sex | EtOH Intake (g/kg/22 h) | Drinking Category | Age (in years) at Start of EtOH Induction | Age Category | Study Inclusion |  |  |  |
| --- | --- | --- | --- | --- | --- | --- | --- | --- | --- | --- |
|  |  |  |  |  |  |  | CAG | TV | BS | WB |
| 10048 | 4 | M | 1.25 | LD | 9.85 | MA | Yes |  | Yes |  |
| 10049 | 4 | M | 2.62 | HD | 9.81 | MA | Yes | Yes | Yes |  |
| 10051 | 4 | M | 2.65 | BD | 9.22 | MA | Yes |  |  |  |
| 10052 | 4 | M | 0.47 | LD | 9.18 | MA | Yes | Yes | Yes |  |
| 10054 | 4 | M | 2.51 | BD | 8.03 | MA | Yes |  |  |  |
| 10055 | 4 | M | 1.03 | LD | 8.43 | MA | Yes | Yes | Yes |  |
| 10056 | 4 | M | 1.56 | LD | 8.01 | MA | Yes | Yes | Yes |  |
| 10057 | 4 | M | 2.29 | BD | 7.05 | MA | Yes |  |  |  |
| 10058 | 4 | M | 1.99 | LD | 7.18 | MA | Yes | Yes |  |  |
| 10059 | 4 | M | 2.43 | BD | 7.18 | MA | Yes |  |  |  |
| 10061 | 5 | M | 3.78 | VHD | 5.85 | YA | Yes | Yes | Yes |  |
| 10062 | 5 | M | 4.09 | VHD | 5.94 | YA |  |  | Yes |  |
| 10063 | 5 | M | 3.09 | VHD | 5.69 | YA |  |  | Yes |  |
| 10064 | 5 | M | 2.74 | HD | 5.65 | YA | Yes |  | Yes |  |
| 10065 | 5 | M | 3.09 | HD | 5.94 | YA | Yes | Yes | Yes |  |
| 10066 | 5 | M | 3.27 | VHD | 5.77 | YA | Yes | Yes | Yes |  |
| 10067 | 5 | M | 3.08 | HD | 5.81 | YA | Yes |  | Yes |  |
| 10167 | 5 | M | - | CTRL | - | - |  |  | Yes |  |
| 10169 | 5 | M | - | CTRL | - | - |  |  | Yes |  |
| 10170 | 5 | M | - | CTRL | - | - |  | Yes | Yes |  |
| 10171 | 5 | M | - | CTRL | - | - |  | Yes | Yes |  |
| 10208 | 10 | M | 2.29 | BD | 5.88 | YA | Yes |  |  |  |
| 10209 | 10 | M | 2.39 | BD | 5.84 | YA | Yes |  |  |  |
| 10210 | 10 | M | 1.55 | LD | 6.16 | YA | Yes |  |  |  |
| 10211 | 10 | M | 2.11 | BD | 5.02 | YA | Yes |  |  |  |
| 10212 | 10 | M | 2.32 | HD | 5.04 | YA | Yes |  |  |  |
| 10213 | 10 | M | 1.29 | LD | 4.74 | Adol | Yes |  |  |  |
| 10214 | 10 | M | 4.22 | VHD | 5.04 | YA | Yes |  |  |  |
| 10215 | 10 | M | 3.19 | VHD | 5.08 | YA | Yes |  |  |  |
| 10068 | 6a | F | n/a | CTRL | - | - |  |  | Yes | Yes |
| 10069 | 6a | F | 5.15 | VHD | 4.14 | Adol | Yes | Yes |  |  |
| 10070 | 6a | F | 3.92 | VHD | 4.12 | Adol | Yes | Yes | Yes |  |
| 10071 | 6a | F | - | CTRL | - | - |  | Yes | Yes | Yes |
| 10076 | 6a | F | - | CTRL | - | - |  | Yes | Yes | Yes |
| 10078 | 6a | F | 5.04 | VHD | 3.84 | Adol | Yes | Yes | Yes | Yes |
| 10079 | 6a | F | 4.02 | VHD | 4.16 | Adol | Yes | Yes | Yes | Yes |
| 10080 | 6a | F | 3.27 | VHD | 4.16 | Adol | Yes | Yes | Yes | Yes |
| 10081 | 6a | F | 4 | VHD | 4.16 | Adol | Yes | Yes |  | Yes |
| 10072 | 6b | F | 1.05 | LD | 5.83 | YA | Yes |  |  | Yes |
| 10073 | 6b | F | 3.94 | VHD | 5.63 | YA | Yes | Yes | Yes |  |
| 10074 | 6b | F | 1.71 | LD | 5.58 | YA | Yes |  |  |  |
| 10075 | 6b | F | 2.78 | HD | 5.83 | YA | Yes |  |  |  |
| 10077 | 6b | F | 1.34 | LD | 6.04 | YA | Yes |  |  |  |
| 10186 | 6b | F | - | CTRL | - | - |  | Yes | Yes | Yes |
| 10187 | 6b | F | - | CTRL | - | - |  | Yes |  | Yes |
| 10188 | 6b | F | - | CTRL | - | - |  | Yes |  |  |
| 10087 | 7a | M | 2.36 | BD | 4.36 | Adol | Yes |  |  |  |

|  |  |  |  |  |  |  |  |  |  |
| --- | --- | --- | --- | --- | --- | --- | --- | --- | --- |
| 10088 | 7a | M | 2.89 | HD | 4.49 | Adol | Yes | Yes |  |
| 10089 | 7a | M | 2 | LD | 4.65 | Adol | Yes | Yes | Yes |
| 10090 | 7a | M | 1.85 | LD | 4.25 | Adol | Yes | Yes | Yes |
| 10091 | 7a | M | 3.15 | VHD | 4.34 | Adol | Yes | Yes |  |
| 10092 | 7a | M | 1.9 | LD | 4.29 | Adol | Yes | Yes | Yes |
| 10093 | 7a | M | - | CTRL | - | - |  | Yes | Yes |
| 10094 | 7a | M | - | CTRL | - | - |  | Yes | Yes |
| 10096 | 7a | M | - | CTRL | - | - |  | Yes | Yes |
| 10097 | 7a | M | 3.03 | HD | 4.33 | Adol | Yes | Yes | Yes |
| 10098 | 7a | M | 3.32 | VHD | 3.99 | Adol | Yes | Yes |  |
| 10082 | 7b | M | 2.42 | HD | 6.30 | YA | Yes | Yes | Yes |
| 10083 | 7b | M | 1.44 | LD | 5.80 | YA | Yes | Yes | Yes |
| 10084 | 7b | M | 1.85 | LD | 5.66 | YA | Yes | Yes | Yes |
| 10085 | 7b | M | 2.07 | LD | 5.87 | YA | Yes | Yes |  |
| 10086 | 7b | M | 2.28 | BD | 5.64 | YA | Yes |  |  |
| 10182 | 7b | M | - | CTRL | - | - |  | Yes | Yes |
| 10183 | 7b | M | - | CTRL | - | - |  | Yes | Yes |
| 10184 | 7b | M | - | CTRL | - | - |  | Yes |  |
| 10185 | 7b | M | - | CTRL | - | - |  | Yes |  |

**Supplemental Table 2.**
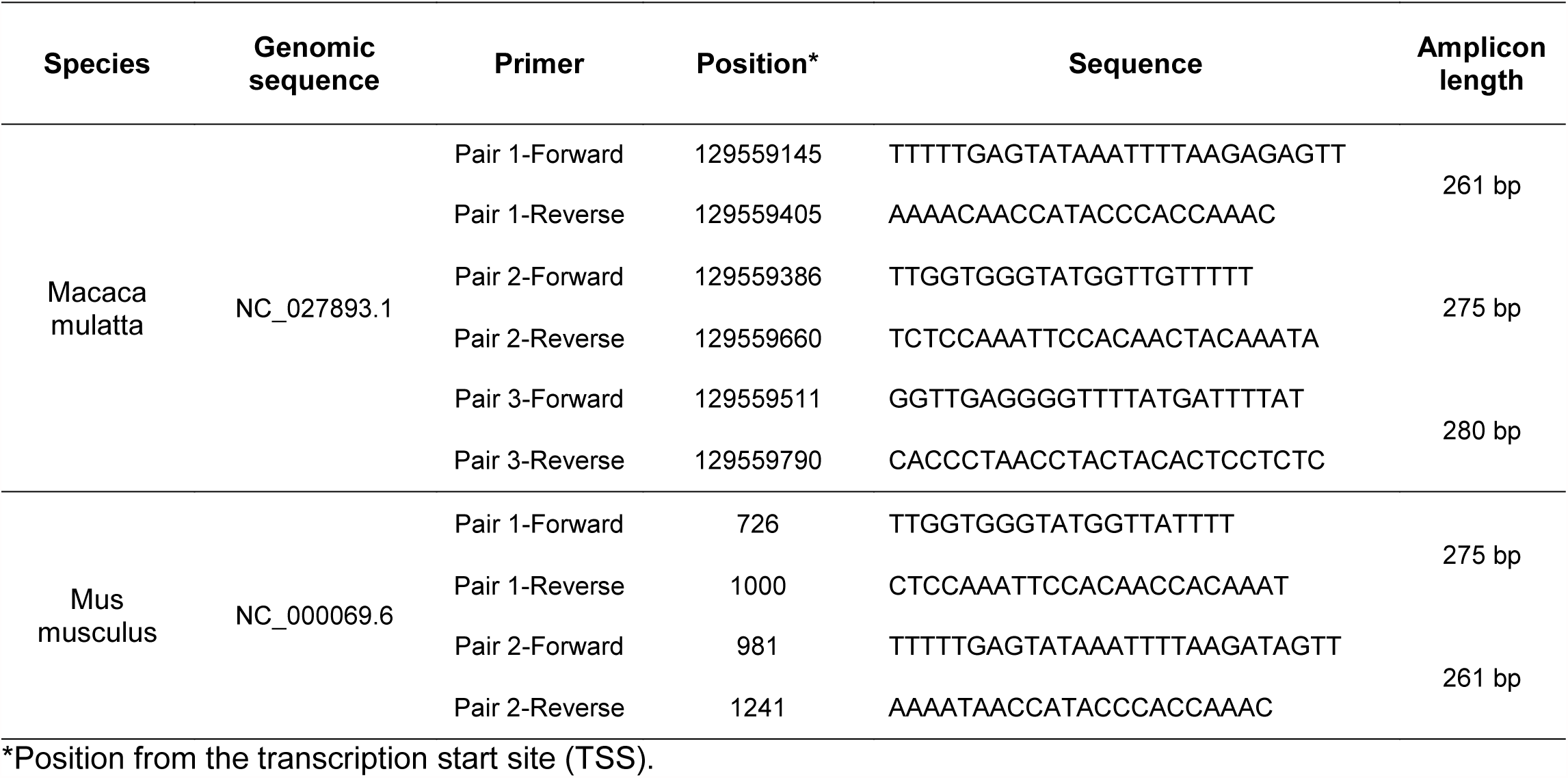
Primers used for bisulfite amplicon sequencing.

**Supplemental Table 3.**
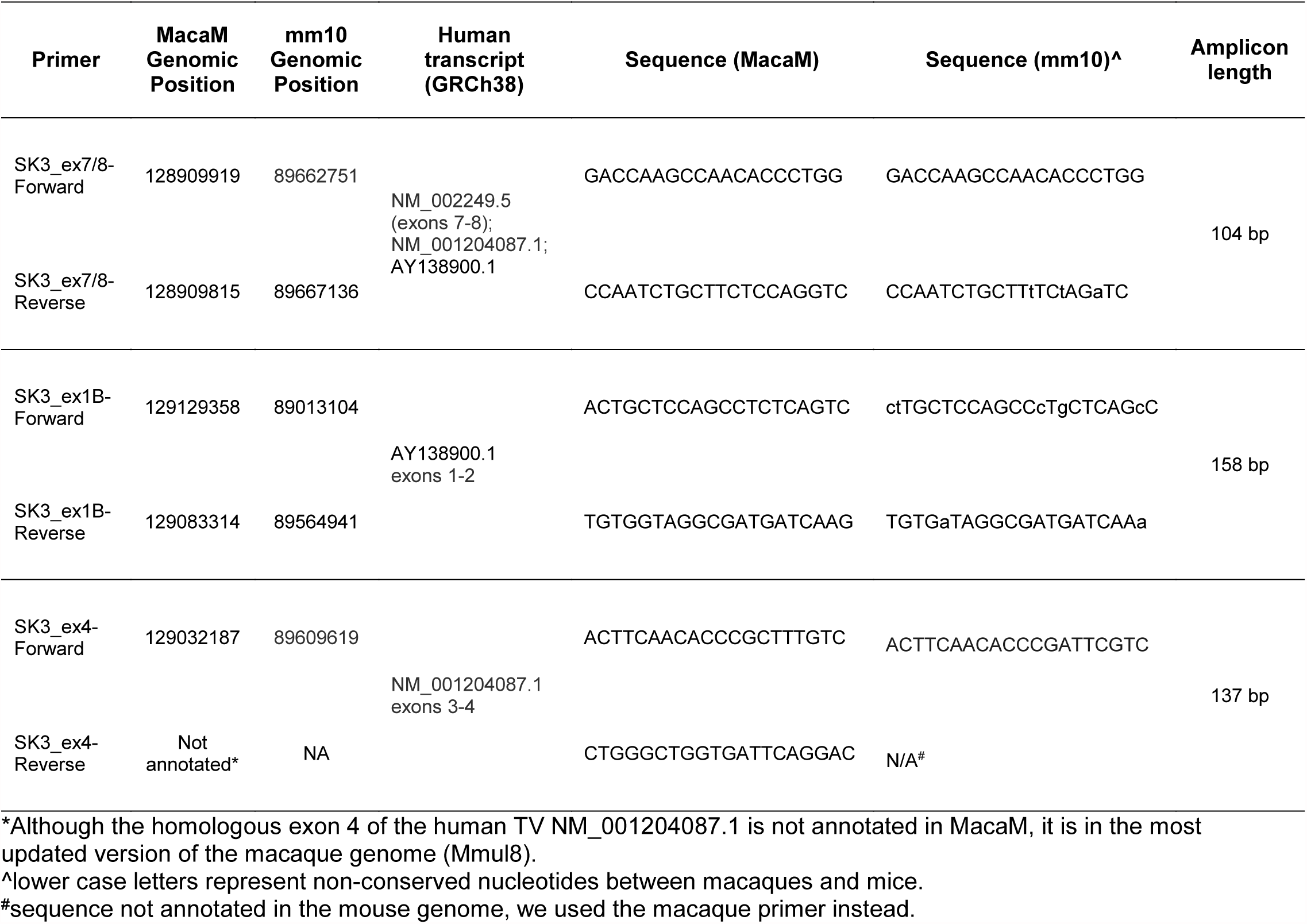
Nucleotide sequences of primers used for quantitative PCR.

**Supplemental Table 4.** *KCNN3* transcript variants, protein isoforms, and channel function.

| Transcript Variant | Unique exons | Protein isoform | Changes in protein structure | Altered Channel Function |
| --- | --- | --- | --- | --- |
| *SK3_1A*<br>(NM_002249.5) | 1A | 1: Full-Length<br>(NP_002240.3) | None | None |
| *SK3_1B*<br>(NM_170782.2) | 1B | 3 (NP_740752.1) | Lacks S1 transmembrane domain; Missing aa1-318 | Dominant-negative |
| *SK3_1C*<br>(AY138900.1) | 1C | 2 (AAN46636.1) | Lacks S1 transmembrane domain; missing aa1-310; 311-315: WGLYS → MERPI | Dominant-negative; Very low CNS expression |
| *SK3_ex4*<br>(NM_001204087.1) | 4 | *SK3_ex4*<br>(NP_001191016) | 15 amino-acid insert in the pore-forming loop | Apamin-insensitive |

**Supplemental Fig. 1.**
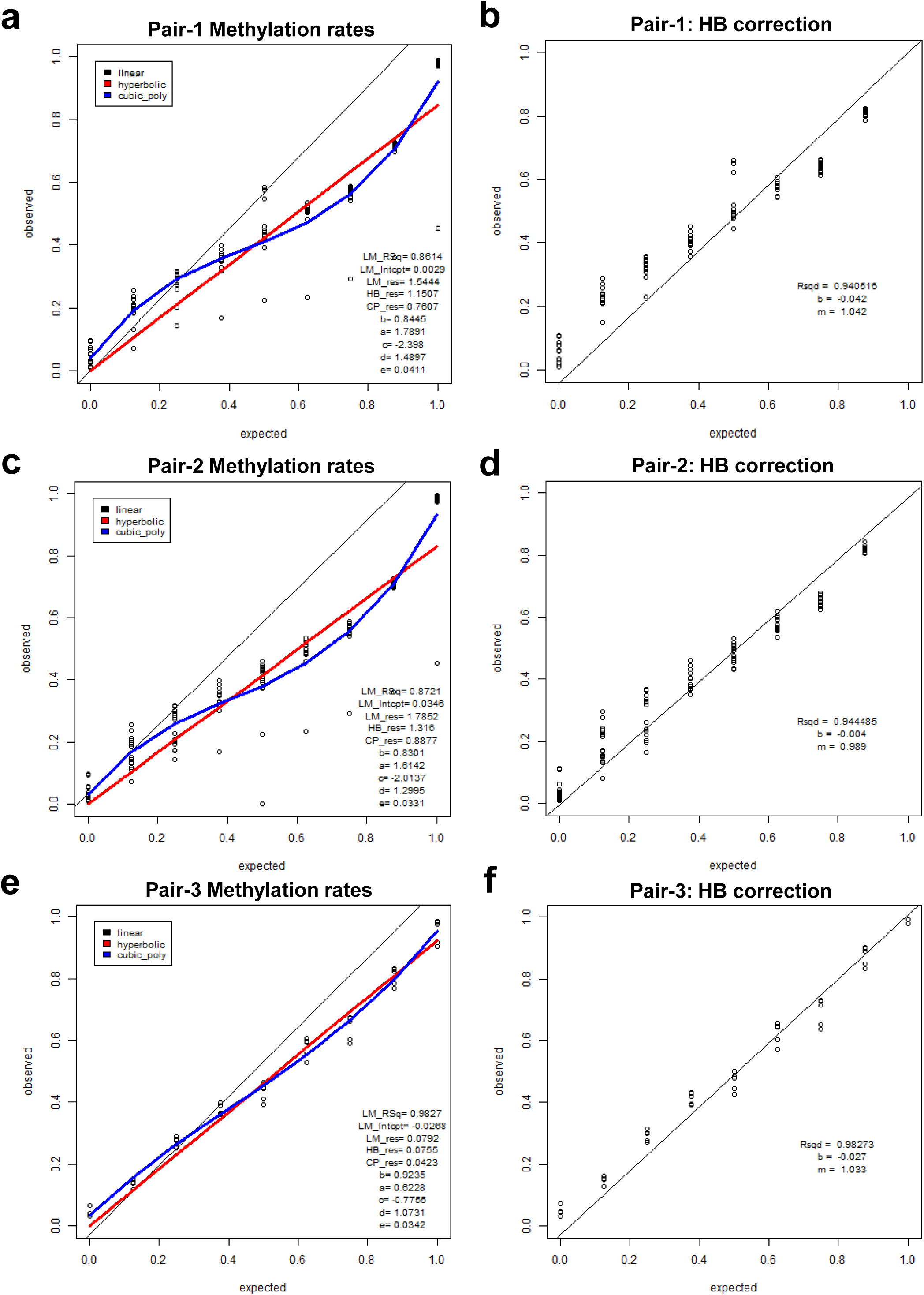
Titration curves to correct for PCR bias during amplification of the methylation region under study. Three different amplicons were used to amplify the complete region 1 (**a**), 2 (**c**) and 3 (**e**). PCR-bias was corrected using the hyperbolic function (HB, **b, d** and **f**).

**Supplemental Fig. 2.**
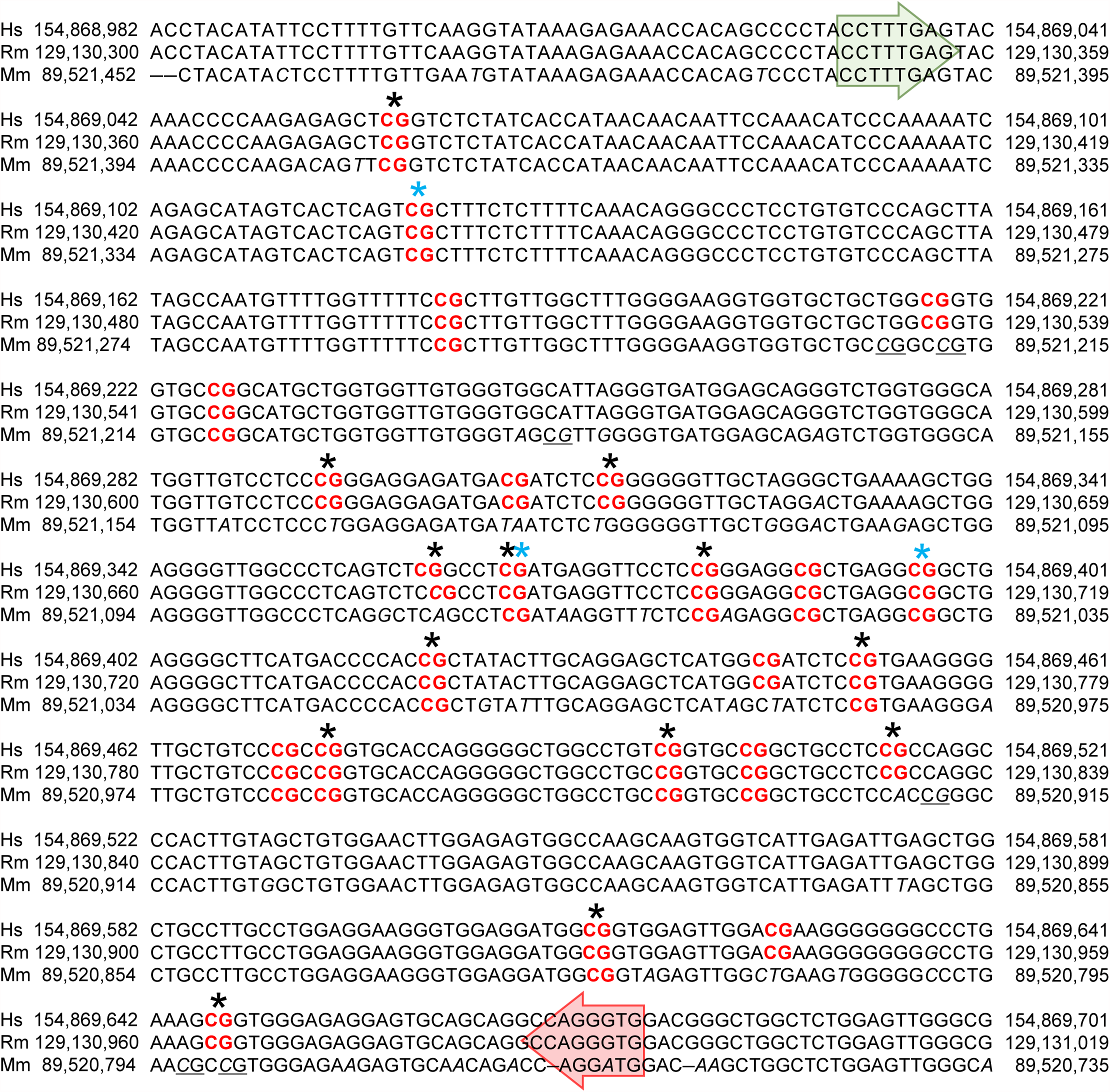
Sequence homology of MR-ex 1 (exon 1/intron 1) across human, macaque and mouse. The start and stop of the differentially methylated region spanning 646 base pairs is shown by green and red arrows, respectively. Conserved CpGs within MR-ex1 are shown in bold, red font. CpGs unique to a species are underlined. Nucleotides that are not conserved across species are shown in italic. ‘*’ and ‘*’ are CpGs that are significantly different in HD/VHD monkeys and CIE-exposed drinking mice, respectively.

**Supplemental Fig. 3.**
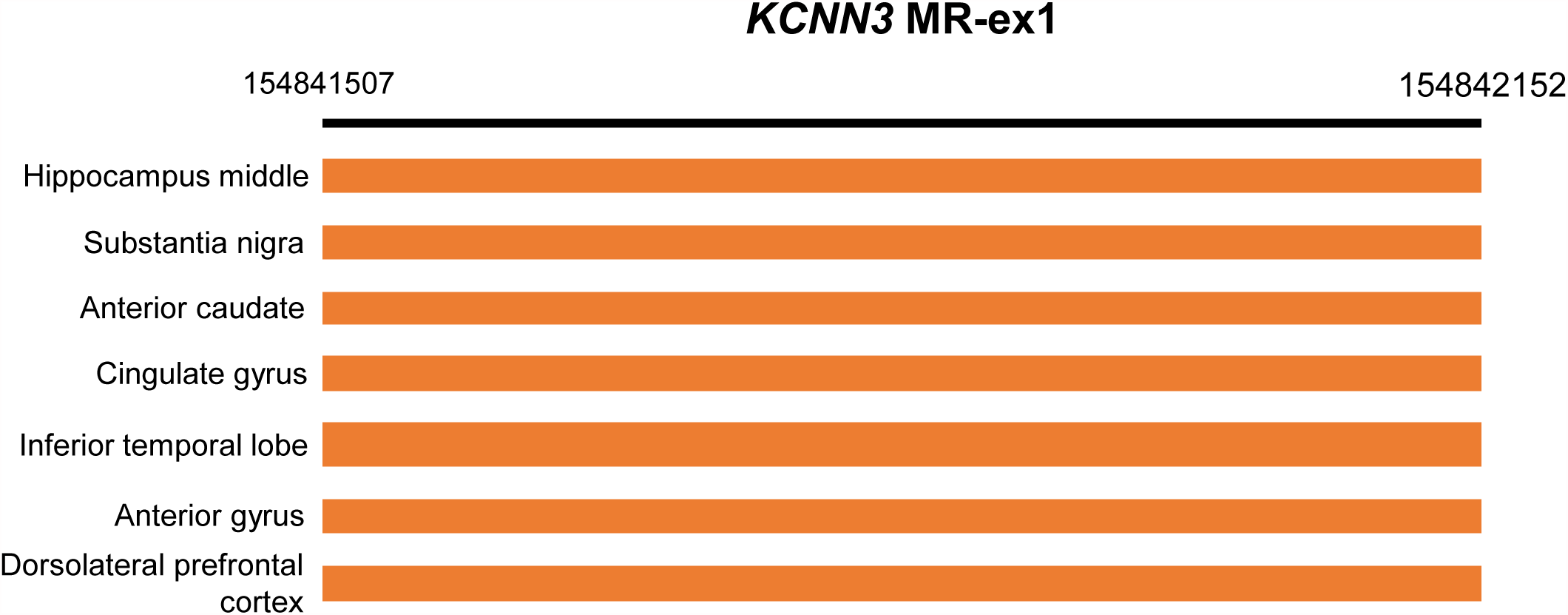
Overlap of the macaque homologous methylation region within MR-ex1 on the human Roadmap 25 chromatin states. The MR-ex1 region on the human genome (hg19) is located at chr1: 154,841,507-154,842,152. According to the Roadmap Epigenomic project, the orange color in the 25 chromatin states represents “promoter upstream transcription start site”. Note that the nucleus accumbens in not included in the Epigenomics Roadmap database.

**Supplemental Fig. 4.**
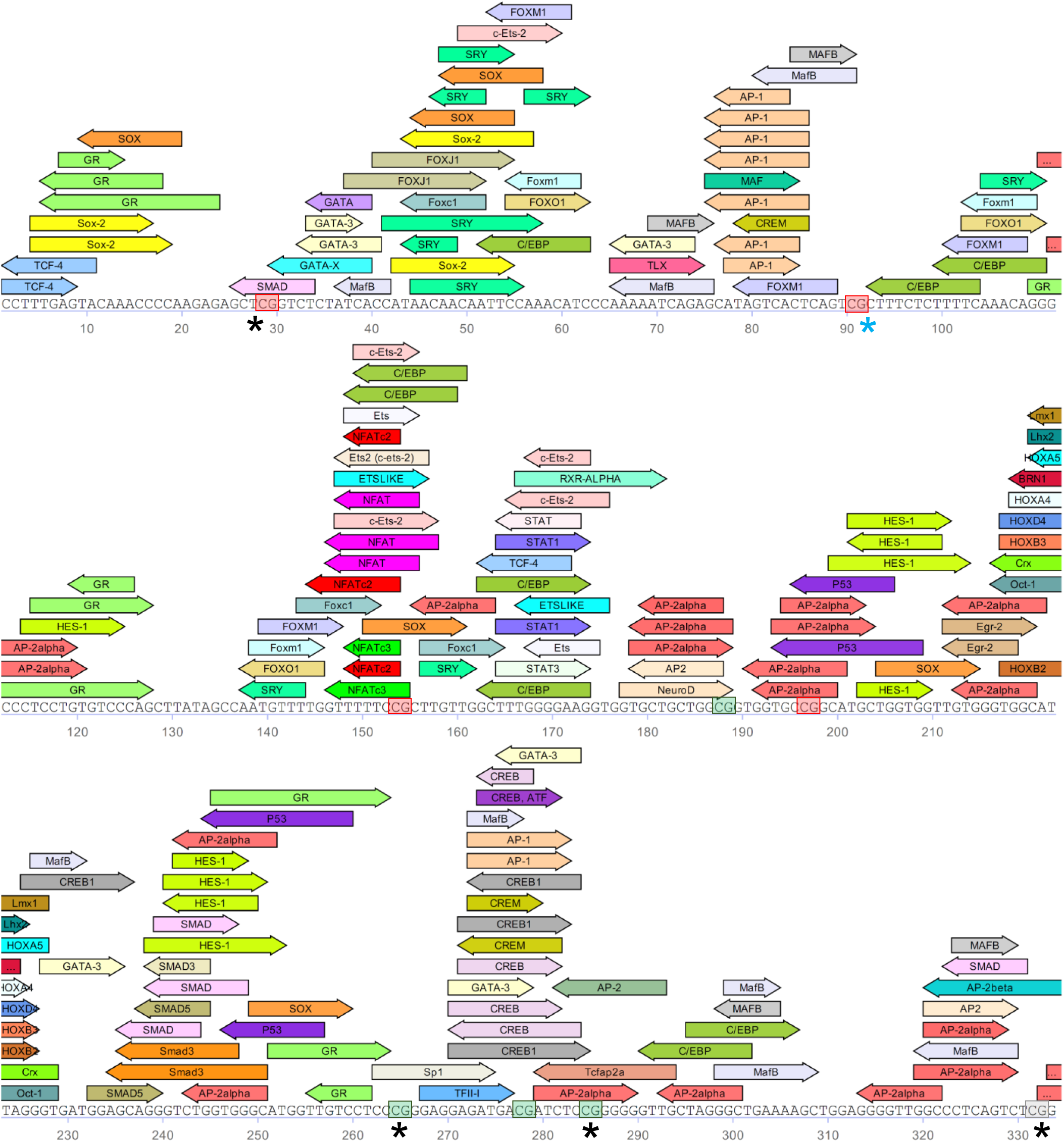
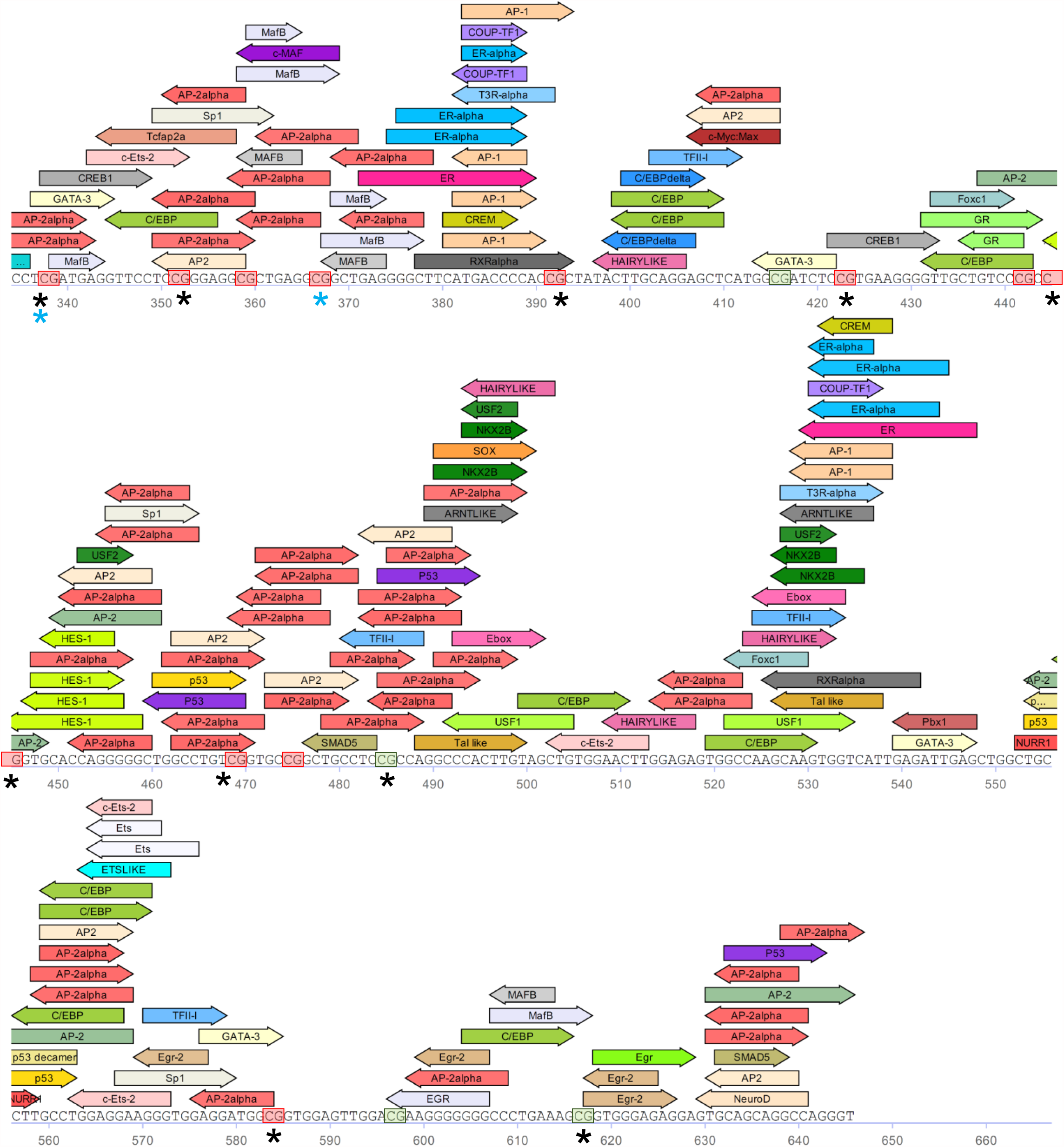
Predicted transcription factors that bind to the human MR-ex1 (TRANSFAC). CpGs enclosed in a red box are conserved in human, monkey and mouse; those CpGs unique to human are enclosed in a grey box, and those conserved in human and monkey are enclosed in a green box. ‘*’ and ‘*’ are CpGs that are significantly different in HD monkeys and CIE-exposed drinking mice, respectively. TRANSFAC conditions were as follows: nerve_system_specific matrices, matrix similarity cut-off: 0.9 and core similarity cut-off: 0.95.

**Supplemental Fig. 5.**
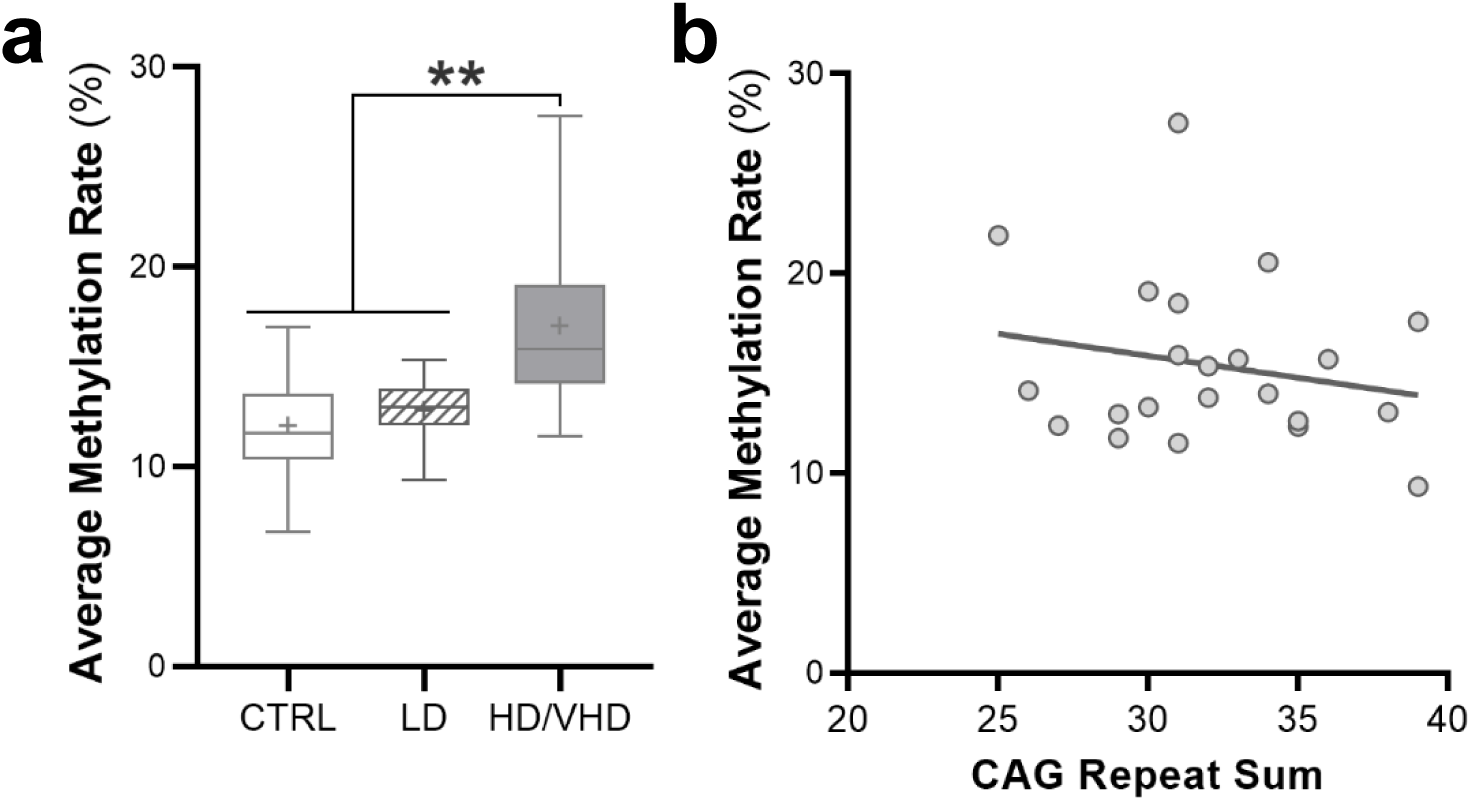
Averaged methylation rate (%) grouped by drinking class for all CpGs in male and female rhesus macaques. **a** The methylation rate was significantly higher in the HD/VHD (N = 15) compared to control (N = 13) and LD (N = 9) macaques (F(2, 34) = 9.860, *p* = 0.0004, Tukey post-hoc, \*\**p* < 0.01). **b** Lack of correlation between average methylation rate and CAG repeat sum (*p* = 0.3563)

